# Chemically-induced epileptic seizures in zebrafish: a systematic review

**DOI:** 10.1101/2023.06.26.546569

**Authors:** Rafael Chitolina, Matheus Gallas-Lopes, Carlos G. Reis, Radharani Benvenutti, Thailana Stahlhofer-Buss, Maria Elisa Calcagnotto, Ana P. Herrmann, Angelo Piato

## Abstract

The use of zebrafish as a model organism is gaining evidence in the field of epilepsy as it may help to understand the mechanisms underlying epileptic seizures. As zebrafish assays became popular, the heterogeneity between protocols increased, making it hard to choose a standard protocol to conduct research while also impairing the comparison of results between studies. We conducted a systematic review to comprehensively profile the chemically-induced seizure models in zebrafish. Literature searches were performed in PubMed, Scopus and Web of Science, followed by a two-step screening process based on inclusion/exclusion criteria. Qualitative data were extracted and a sample of 100 studies was randomly selected for risk of bias assessment. Out of the 1058 studies identified after removing duplicates, 201 met the inclusion criteria. We found that the most common chemoconvulsants used in the reviewed studies were pentylenetetrazole (n = 180), kainic acid (n = 11), and pilocarpine (n = 10), which increase seizure severity in a dose-dependent manner. The main outcomes assessed were seizure scores and locomotion. Significant variability between the protocols was observed for administration route, duration of exposure, and dose/concentration. Of the studies subjected to risk of bias assessment, most were rated as low risk of bias for selective reporting (94%), baseline characteristics of the animals (67%), and blinded outcome assessment (54%). Randomization procedures and incomplete data were rated as unclear in 81% and 68% of the studies, respectively. None of the studies reported the sample size calculation. Overall, these findings underscore the need for improved methodological and reporting practices to enhance reproducibility and reliability of zebrafish models for studying epilepsy. Our study offers a comprehensive overview of the current state of chemically-induced seizure models in zebrafish, highlighting the common chemoconvulsants used and the variability in protocol parameters. This may be particularly valuable to researchers interested in understanding the underlying mechanisms of epileptic seizures and screening potential drug candidates in zebrafish models.

**HIGHLIGHTS:**

- We systematically reviewed the effects of chemically-induced seizures in zebrafish;
- PTZ is the most used epileptic seizure inducer in zebrafish;
- More than 50% of the studies fail to report data such as outlier exclusion criteria and sample size estimation;
- The results showed a need for better standardization of protocols.

## INTRODUCTION

Among the main chronic neurological diseases, epilepsy is a condition characterized by the occurrence of recurrent unprovoked epileptic seizures that affects around 70 million people worldwide [1,2]. Epilepsy is defined by the International League Against Epilepsy (ILAE) as a “disorder of the brain characterized by an enduring predisposition to generate epileptic seizures and by the neurobiologic, cognitive, psychological, and social consequences of this condition” [1]. An epileptic seizure is defined as a transient occurrence of signs and/or symptoms due to abnormal or excessive neuronal activity in the brain, and is characterized by stereotyped behavioral changes that reflect the underlying neural mechanisms of the disease [1,3]. Epileptogenesis is triggered by various pathogenetic events, encompassing acquired causes (e.g. stroke or traumatic brain injury) or infectious diseases, autoimmune diseases, or a genetic alteration, However, the etiology remains unidentified in a significant number of patients. This process commences prior to and continues beyond the occurrence of the initial unprovoked seizure. [4]. To date, more than 500 genes associated with epilepsy have been identified [4].

Patients with epilepsy are mainly treated with antiseizure drugs (ASDs), aiming to reduce or prevent the occurrence of seizures. After starting the treatment with an antiseizure medication, 80% of patients experience adverse effects that impair their quality of life or result in interruption or non-adherence to treatment [4]. Also, despite the more than 20 antiepileptic drugs currently available, approximately 30 to 40% of patients present with refractory epilepsy, when seizures are not controlled by two or more antiseizure drugs [5]. To address this issue, there is a pressing need to develop novel treatment approaches and antiepileptic drugs for combating epilepsy [6].

Most of the studies in the literature involving the process of epileptogenesis are carried out in rodent models [4]. Zebrafish (*Danio rerio* Hamilton, 1822) is a model organism that has been gaining more space in the field of epileptogenesis, behavior during epileptic crises, and in the discovery of new antiseizure drugs and therapeutic targets [7–9]. Zebrafish has been widely used in neuroscience research as it is a vertebrate species with a central nervous system architecture similar to mammals and physiological and genetic properties homologous to humans [10]. In addition, because of its external development, zebrafish present advantages in terms of genetic manipulation techniques when compared to rodents [11]. Also, zebrafish can absorb compounds added to water and have a characterized and detailed behavioral pattern [10]. Studies have shown that 71% of the genes encoding proteins in the human genome are related to genes found in the zebrafish genome, and 84% of the genes associated with human diseases have a zebrafish homolog, proving to be a useful model organism also for biochemical and genetic studies [12].

Different behavioral assays of epileptic seizures in zebrafish have been described; seizure-inducing agents, such as pentylenetetrazole, picrotoxin, and allylglycine [7,13,14], among others, are employed in varying concentrations and routes of administration. These behavioral studies have been of great importance in the discovery of new compounds or potential antiseizure drug candidates capable of reducing the stages of epileptic seizures that mimic the crises that happen in humans. However, the heterogeneity in terms of methodologies described in the published literature is high, where researchers tend to adapt protocols (e.g., use a different time of exposure to a seizure inducer, or a different concentration). In addition, there are differences in the behavioral characterization of the effects during exposure to epileptic seizure inducers.

We conducted a systematic review of studies using chemical inducers of epileptic seizures in zebrafish available in the indexed literature. In addition to their use in clinical research, systematic reviews are a tool used to externally validate models and protocols used in preclinical research. They contribute to improve these methods by the investigation of potential sources of heterogeneity and biases, also indicating the quality of the studies. We qualitatively describe the published studies, annotating the results on neurobehavioural and neurochemical parameters, detecting patterns and possible effect moderators, and evaluating the impact of bias arising from the methodological conduct and reporting quality.

## METHODS

This review followed our predefined SYRCLE (Systematic Review Center for Laboratory Animal Experimentation) protocol [15] registered in the Open Science Framework prior to the screening of records and data collection. Preregistration is available at https://osf.io/2njhw [16]. The reporting of this study complies with the Preferred Reporting Items for Systematic Reviews (PRISMA) guidelines [17].

### Search Strategy

Studies were identified through a scientific literature search in three different databases: PubMed, SCOPUS, and Web of Science. Search strategies were designed to suit the characteristics of each database. Varied terms were used for the searches that describe the intervention (epileptic seizure induction protocol) combined with terms referring to the population of interest (zebrafish), in order to carry out a sensitive search. The complete query for each database can be found at https://osf.io/2njhw [16]. The searches were performed without restrictions for language or year of publication, on November 5^th^, 2021. The bibliographic data acquired were imported into Rayyan software [18], where duplicates were detected and removed by one of the investigators (RC). The reference lists of the included studies were also screened to detect additional relevant articles.

### Eligibility screening

After the removal of duplicates, the selection of eligible studies was conducted using Rayyan software. The records underwent a pre-selection based on their title and abstract. For records that did not fit the determined exclusion criteria, an analysis of the full text was carried out. In both stages (title/abstract and full-text analysis) two independent reviewers analyzed each study (RC and MGL or RB or CGR). Disagreements between reviewers’ decisions were resolved by a third reviewer (MEC or APH or AP). Experimental studies using chemicals to induce seizures in zebrafish on the following outcomes were included: behavioral parameters during epileptic seizures (distance traveled, immobility, frequency, and duration of epileptic seizures, latency for stages of epileptic seizures), neurochemical outcomes (e.g., neurotransmitter levels, gene expression, protein levels, oxidative stress), and late behavioral effects (e.g., outcomes related to cognition and social behavior).

In the first screening stage (title and abstract), studies were excluded based on the following reasons: (1) design: not an original primary study (e.g., review, commentary, conference proceedings, and corrections); (2) population: studies performed in animals other than zebrafish (*Danio rerio*) or *in vitro*, *ex vivo* studies; (3) intervention: other methods of seizure induction than chemically-induced (e.g., genetically modified animals, electroshock). In the second stage (full-text screening), the remaining articles were assessed for exclusion based on the same reasons considered in the first stage plus the following additional reasons: (4) outcome: studies that did not assess behavioral or neurochemical parameters.

### Data extraction

Data extraction from the included studies was conducted by two independent investigators (RC and MGL or RB or CGR or TSB) and disagreements were resolved by a discussion between the two investigators. Whenever available, the exact information was extracted directly from text or table. When information was not found, we used unclear.

The following characteristics were extracted: (1) study characteristics: study title, digital object identifier (DOI), first and last authors, last author’s institutional affiliation, and year of publication; (2) animal model characteristics: strain, sex, developmental stage during exposure to the chemical inducer; (3) protocol characteristics: seizure inducer, administration route, frequency of exposure, duration of exposure, dose or concentration, interval between inducer exposure and test, antiepileptic treatment; (4) test characteristics: type of test, test duration, category of measured variable, measured variable and main findings of the study.

To assess the relationship between researchers that publish within this field, co authorship networks were constructed using VOSviewer software version 1.6.18 (https://www.vosviewer.com) [19,20].

### Risk of bias and reporting quality

To evaluate the quality of the included studies, a sample of 100 articles was chosen at random (using random.org) for risk of bias analysis. The risk of bias assessment was conducted by two independent investigators for each paper (RC and MGL or RB or CGR or TSB), and disagreements were resolved by discussion between the two investigators. The analysis was conducted based on the SYRCLE’s risk of bias tool for animal studies [21], with adaptations to better suit the model animal and the intervention of interest. The following items were evaluated for methodological quality: (1) description of random allocation of animals; (2) description of baseline characteristics; (3) description of blinding methods for outcome assessment; (4) incomplete outcome data; (5) selective outcome reporting. Additionally, two other items were evaluated by the investigators to assess the overall reporting quality of the studies based on a set of reporting standards [22]: (6.1) sample size estimation; (6.2) mention of inclusion/exclusion of samples criteria. For methodological quality, each item was scored with a “Yes” for low risk of bias, “No” for a high risk of bias, or “Unclear” when it was not possible to estimate the risk of bias based on the information provided. Items regarding reporting quality were scored with only “Yes„ or “No„, meaning low or high methodological reporting quality, respectively. A complete guide for assessing the risk of bias associated with each of the items in this review is available at https://osf.io/z698v. Risk of bias plots were created using *robvis* [23].

## RESULTS

### Search results

From the search in the selected databases, 2130 records were retrieved altogether (Pubmed = 807; Scopus = 710; Web of Science = 613). Following the removal of duplicates, 1.058 records were screened for eligibility based on title and abstract. After the first screening phase, 283 studies remained to be assessed based on full-text, and 201 met the criteria and were included in the review (Fig. 1). In the two screening phases (title/abstract and full text), 299 did not use a chemical inducer of epileptic seizures as an intervention, 277 articles were excluded due to the study design, 268 did not meet the population criteria and 10 did not report any of the outcomes of interest. In addition, 3 studies were not retrieved and were therefore excluded from the final analysis. This resulted in a total of 201 studies being included in the qualitative synthesis. The flowchart illustrates the progressive selection of studies and the number of articles in each stage (Fig. 1).

**Fig. 1.**
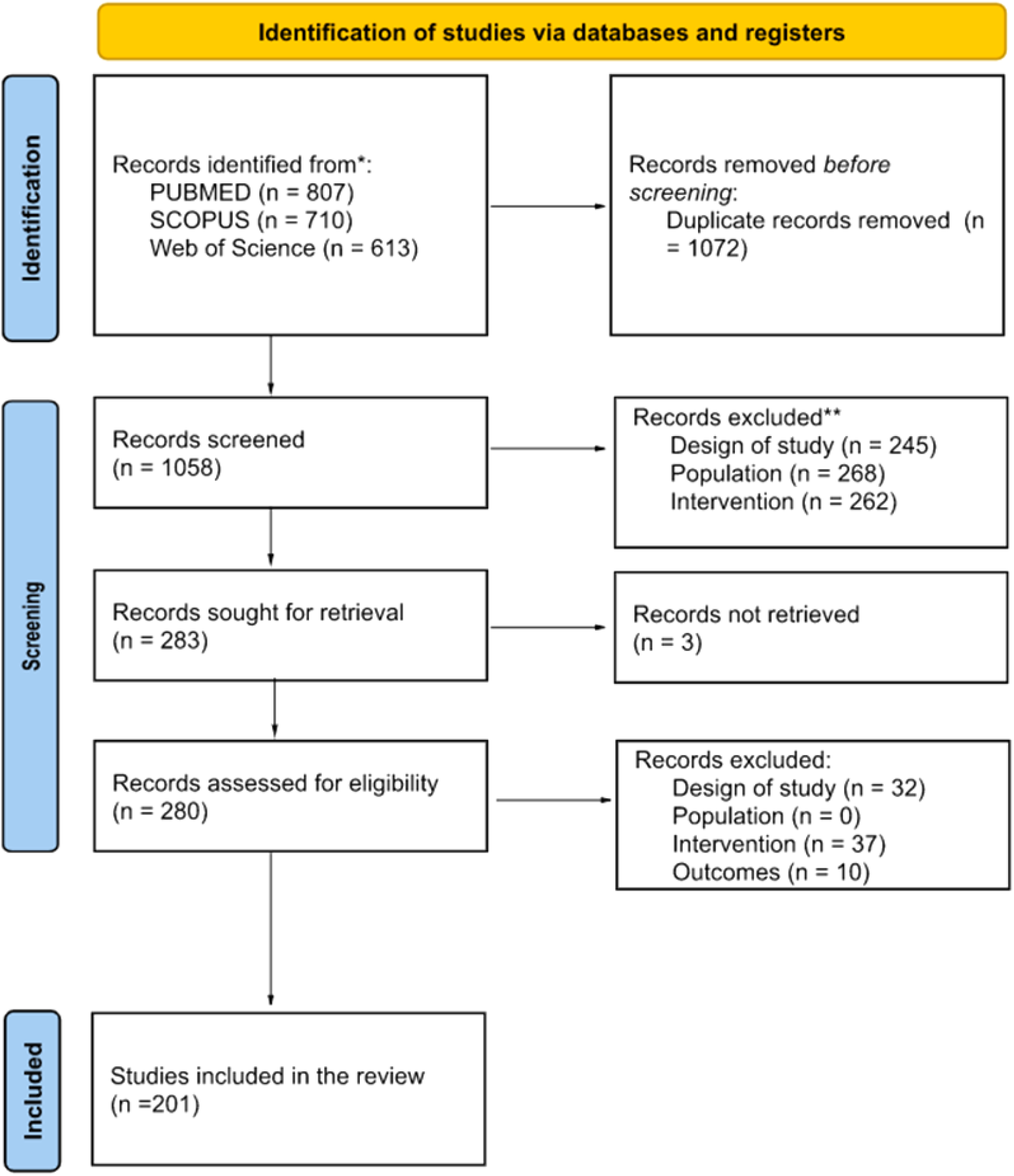
Flowchart diagram of the collection of studies and selection process of the systematic review according to the PRISMA statement.

### Study characteristics

In the 201 studies included in this review, most used pentylenetetrazole (PTZ) as an epileptic seizure inducer (n = 180, 89.55%); other inducers used to study epilepsy were kainic acid (n = 11, 5.47%), pilocarpine (PILO) (n = 8, 3.98%), picrotoxin (PTX) (n = 8, 3.98%), domoic acid (DA) (n = 5, 2.48%), 4-aminopyridine (4-AP) (n = 5, 2.48%), caffeine (n = 3, 1.49%) and ethyl ketopentenoate (EKP) (n = 3, 1.49%). In addition to the main inducers found, others appeared once or twice, namely: strychnine, ginkgotoxin (GT), allylglycine (AG), triocresyl-phosphate (TCP), 1,3,5-trinitroperhydro-1,3,5-triazine (RDX), N-methyl-D-aspartic acid (NMDA) and aconitine. It is important to mention that, in some studies, more than one seizure inducer was used, as described in Table 1.

**Table 1.**
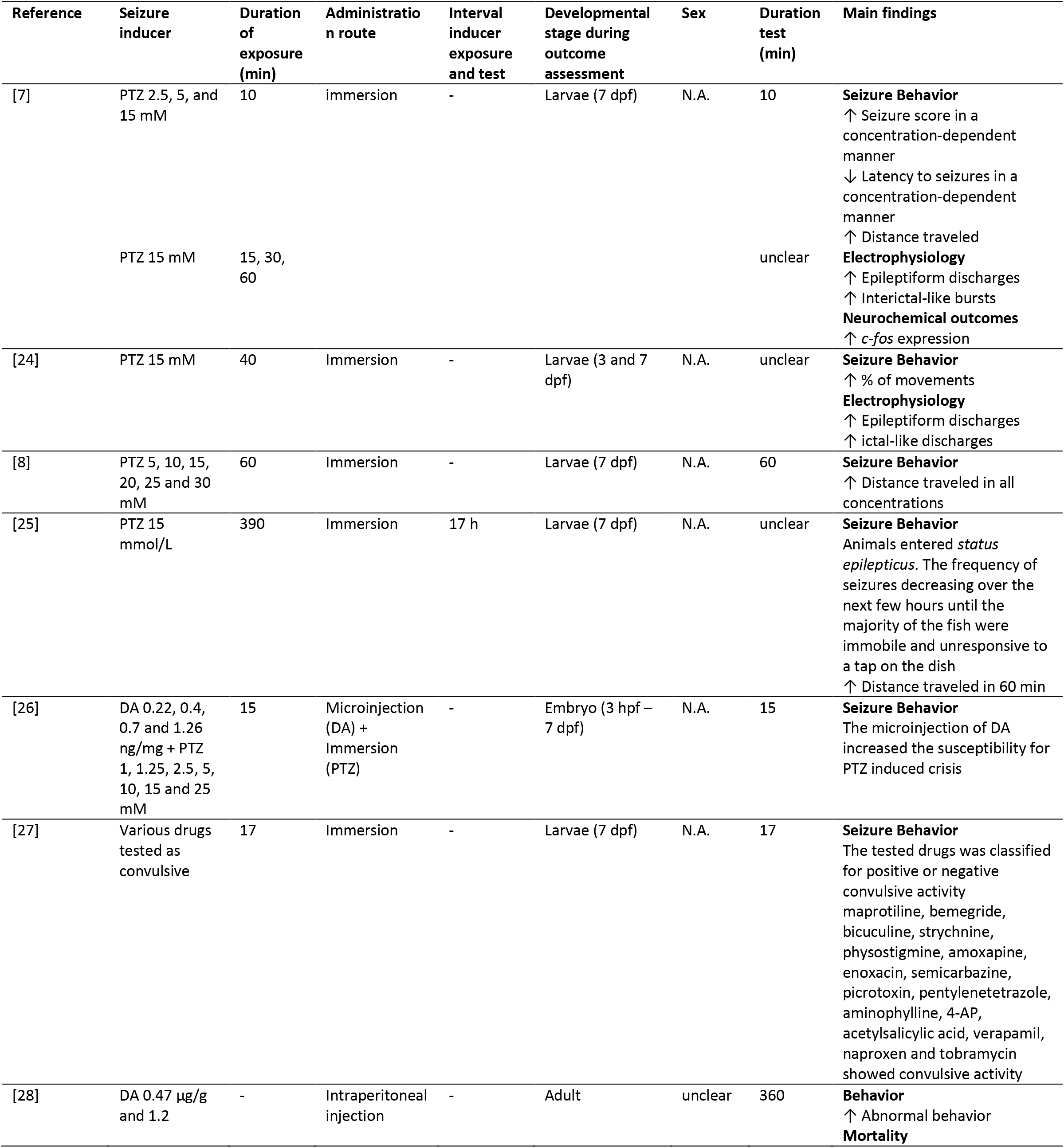
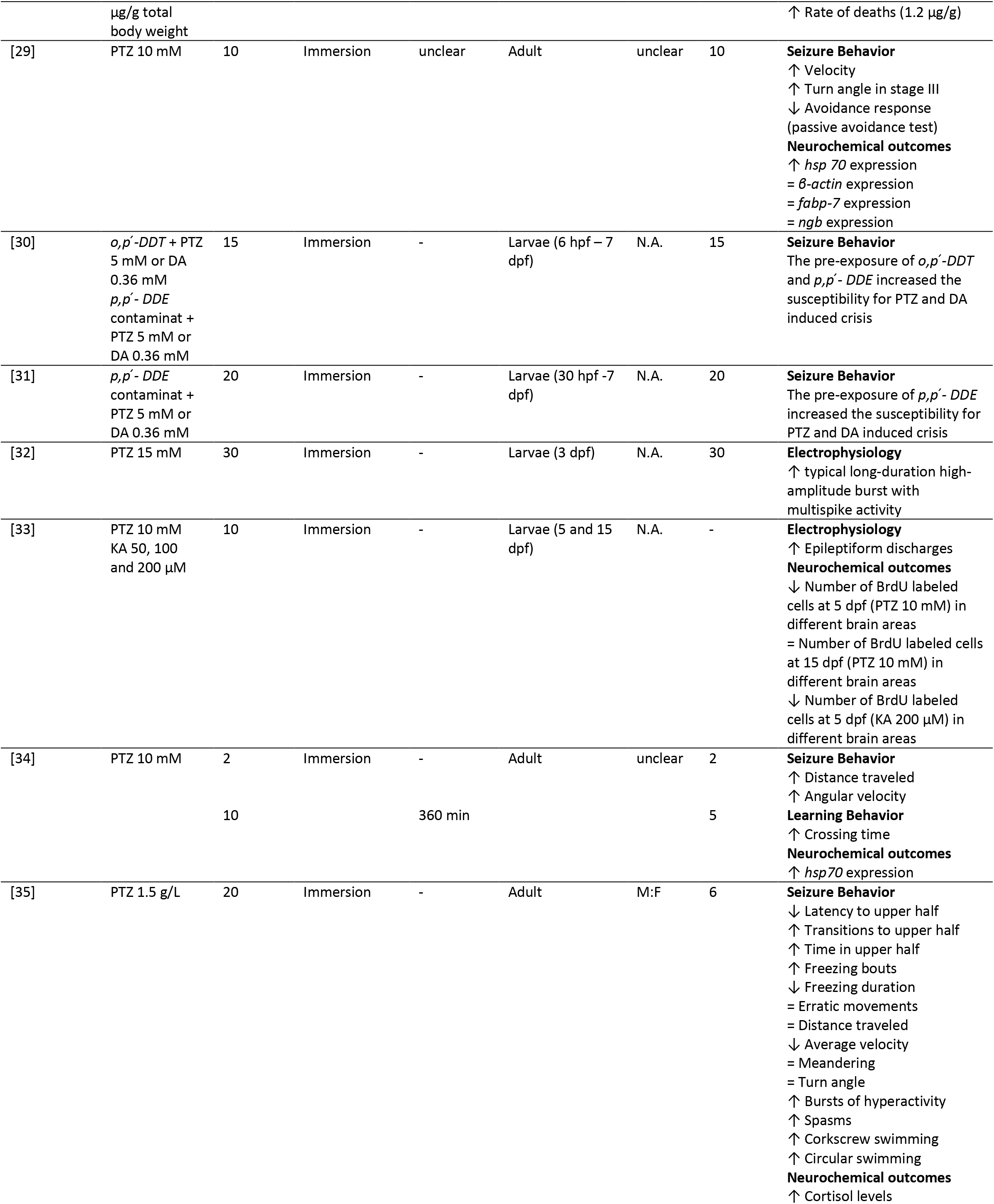
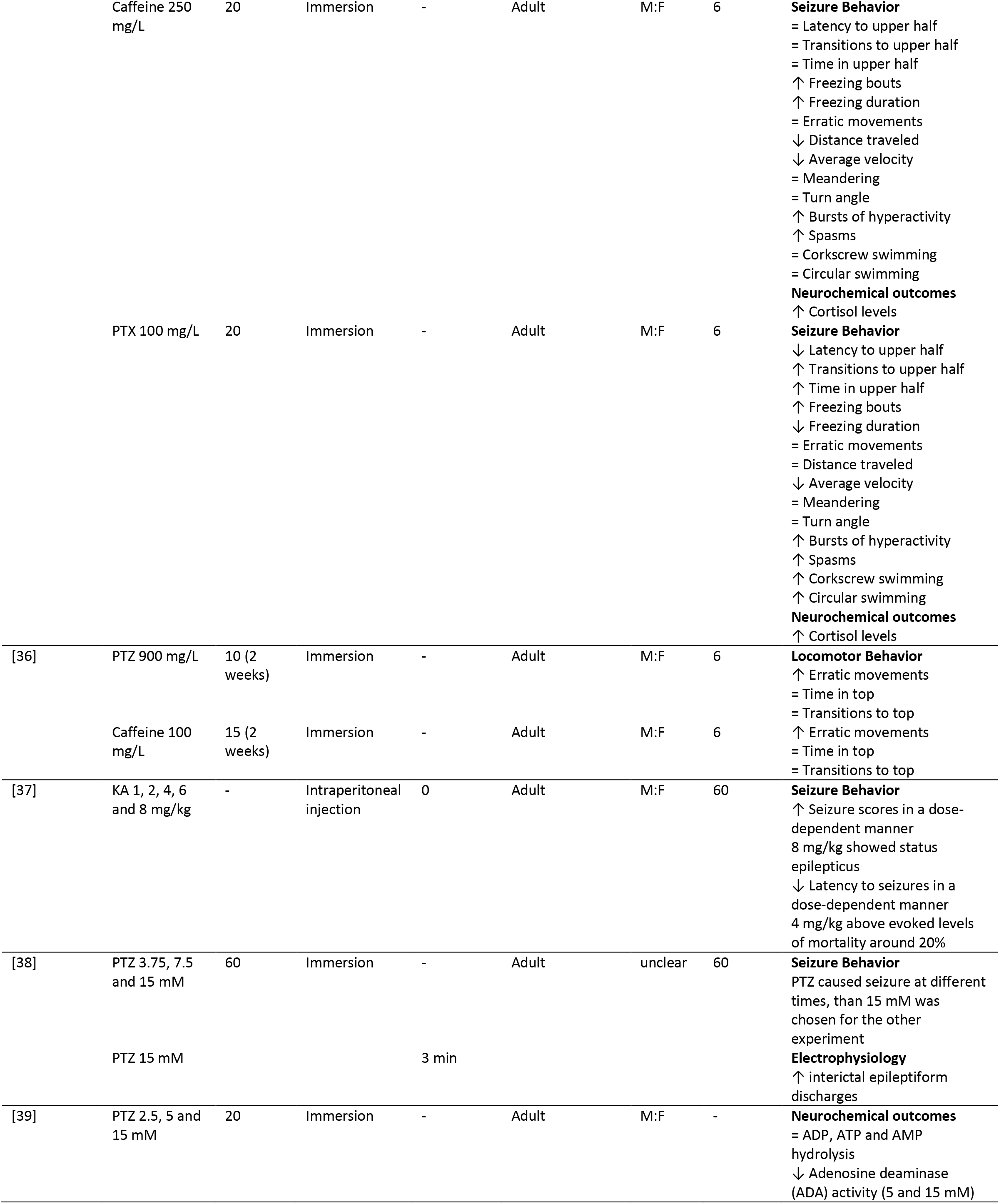
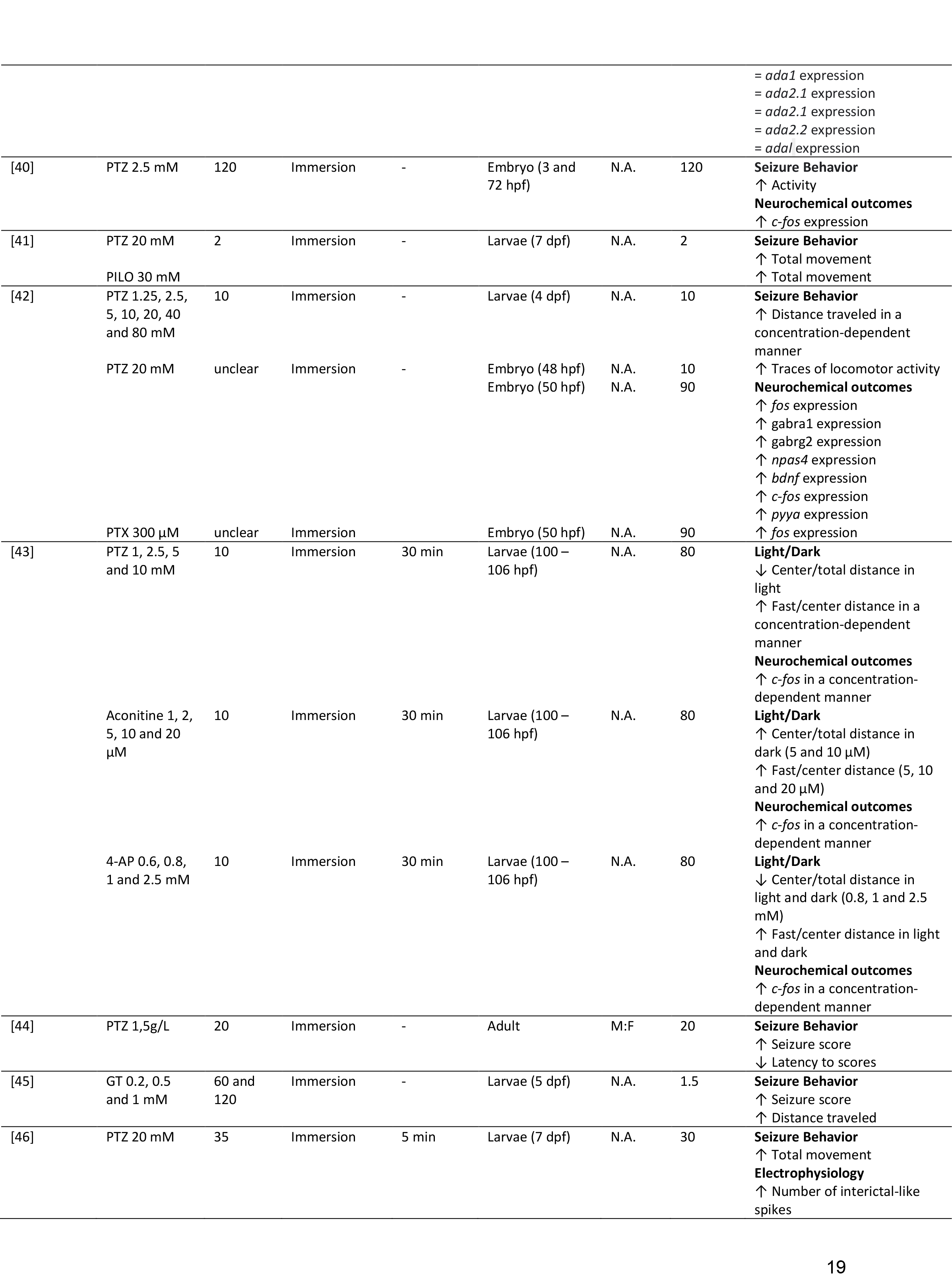
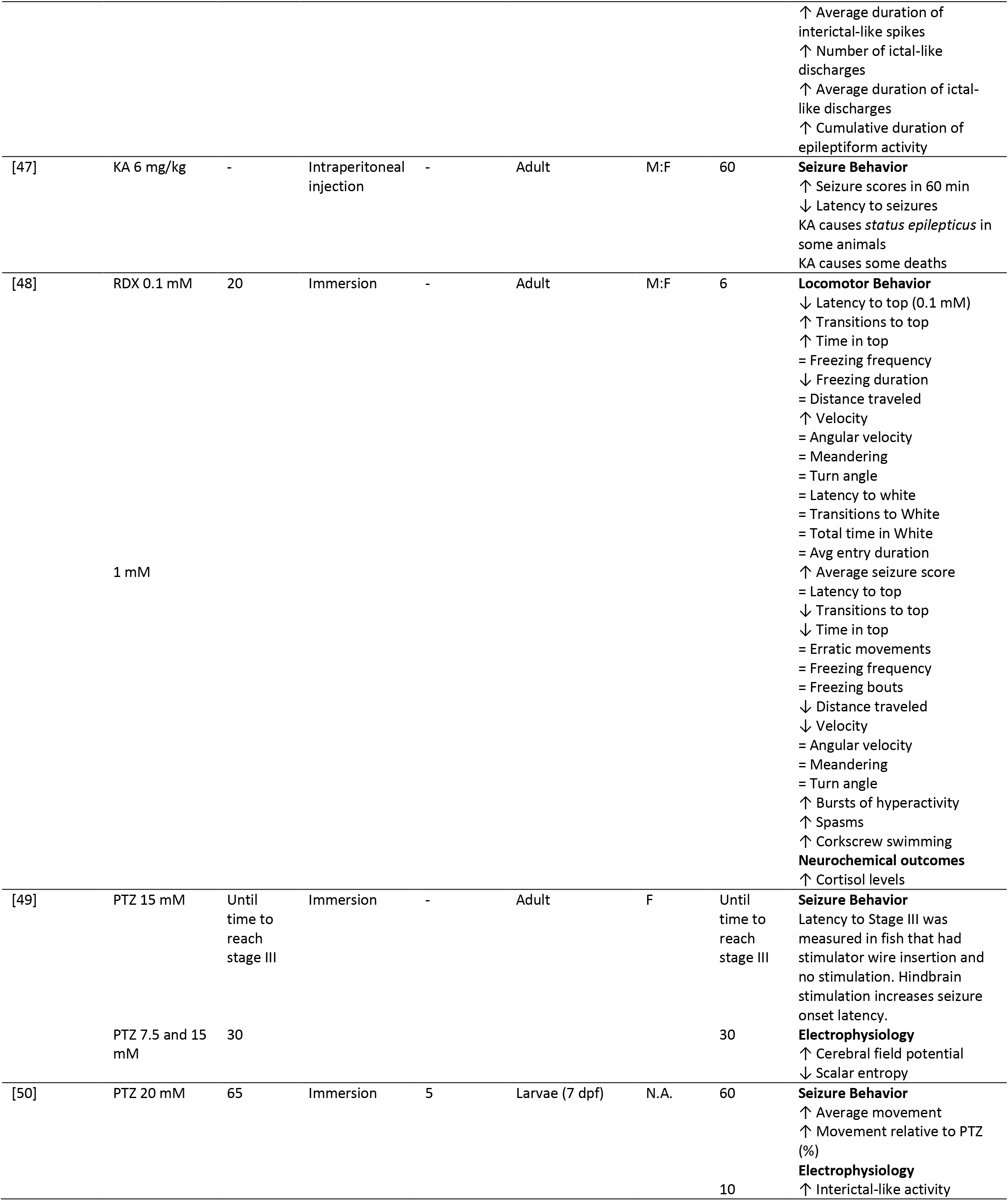
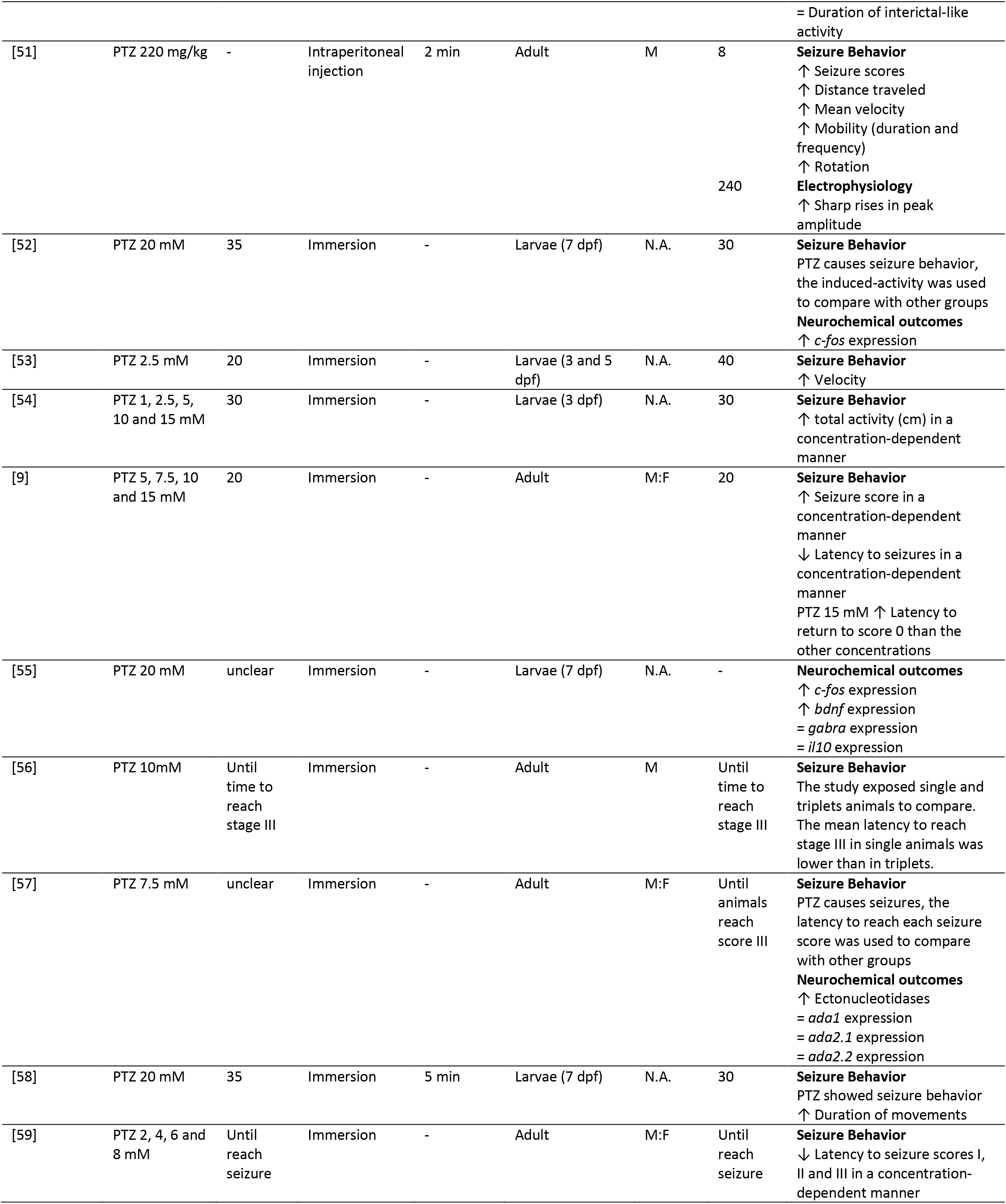
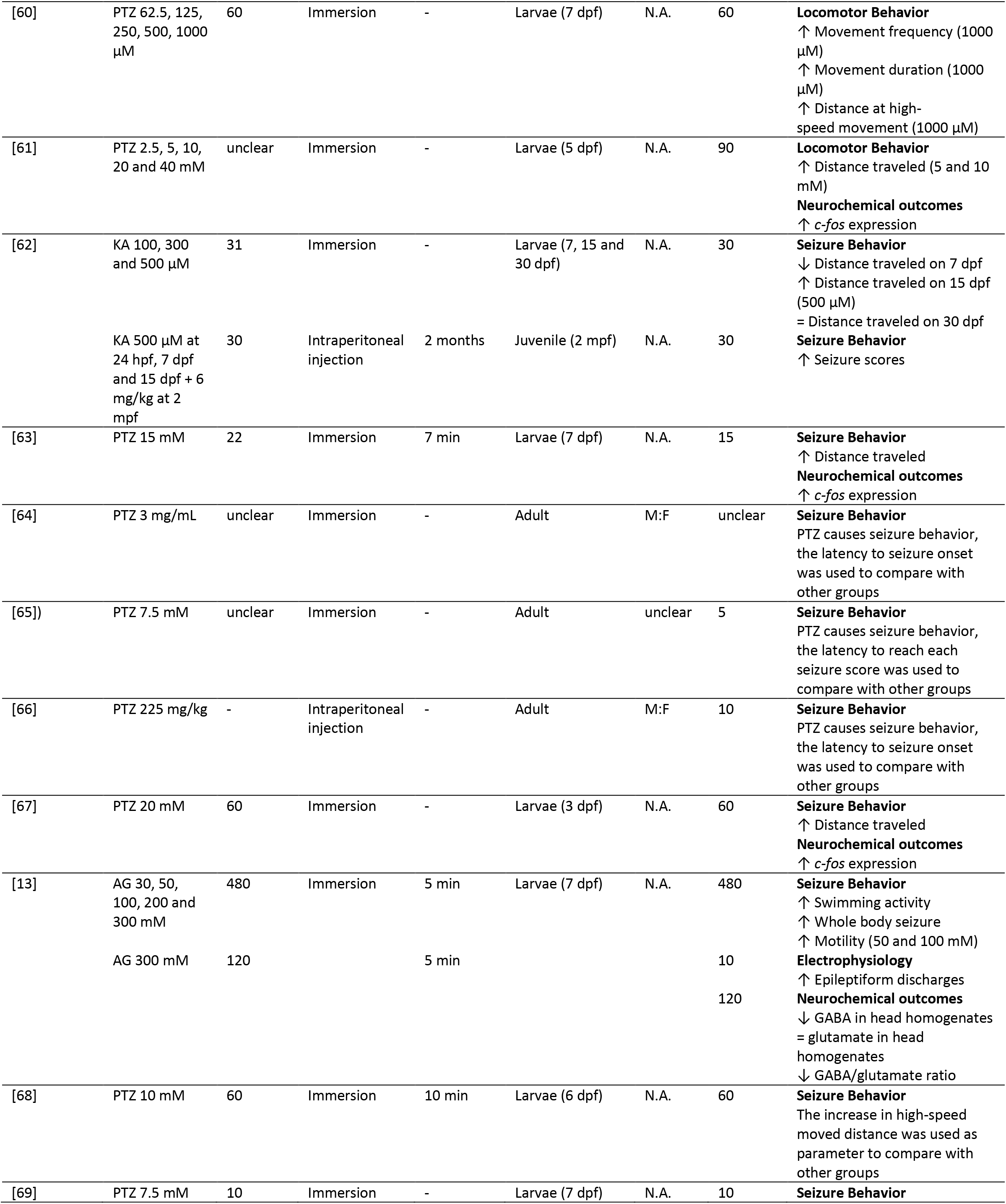
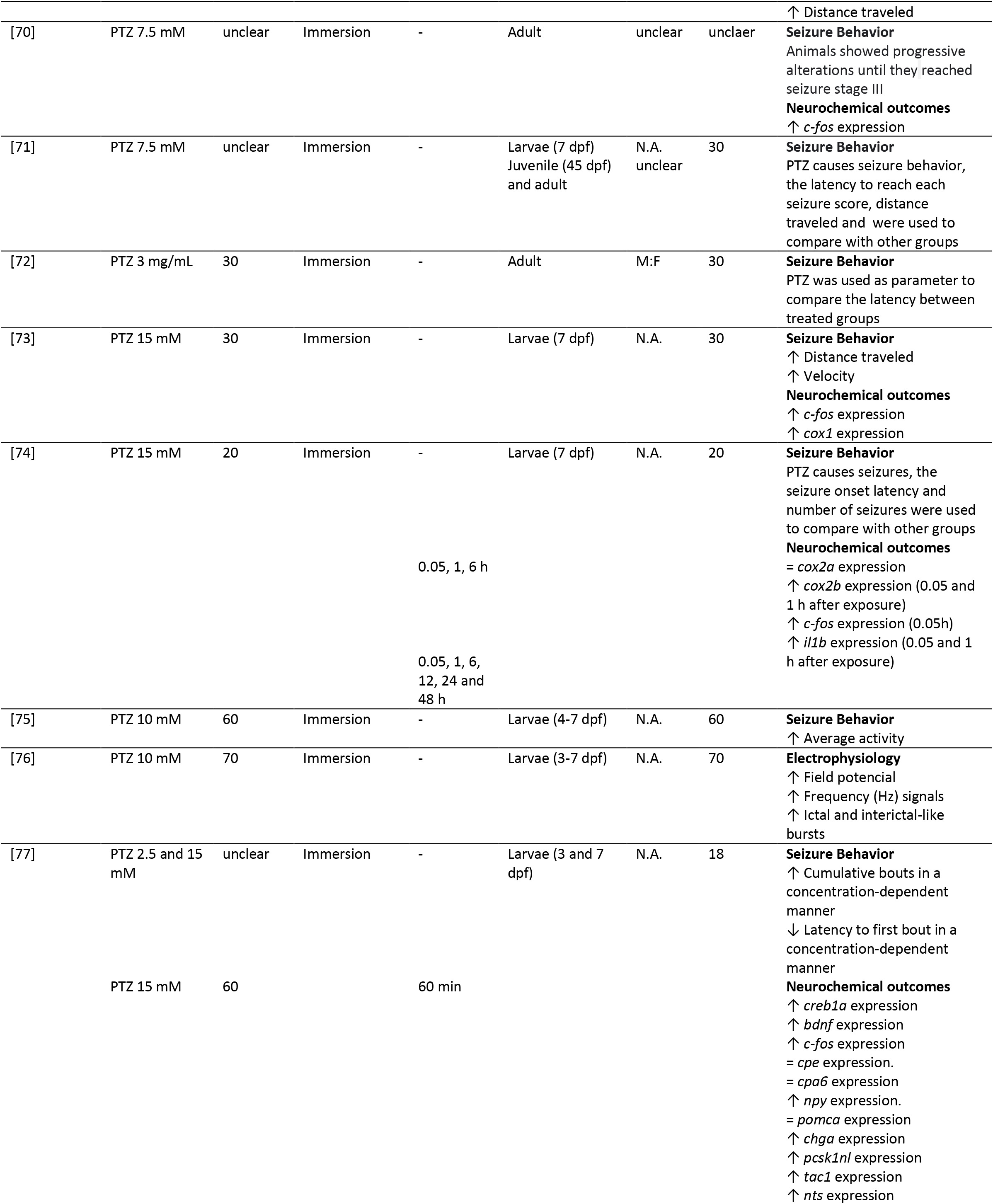
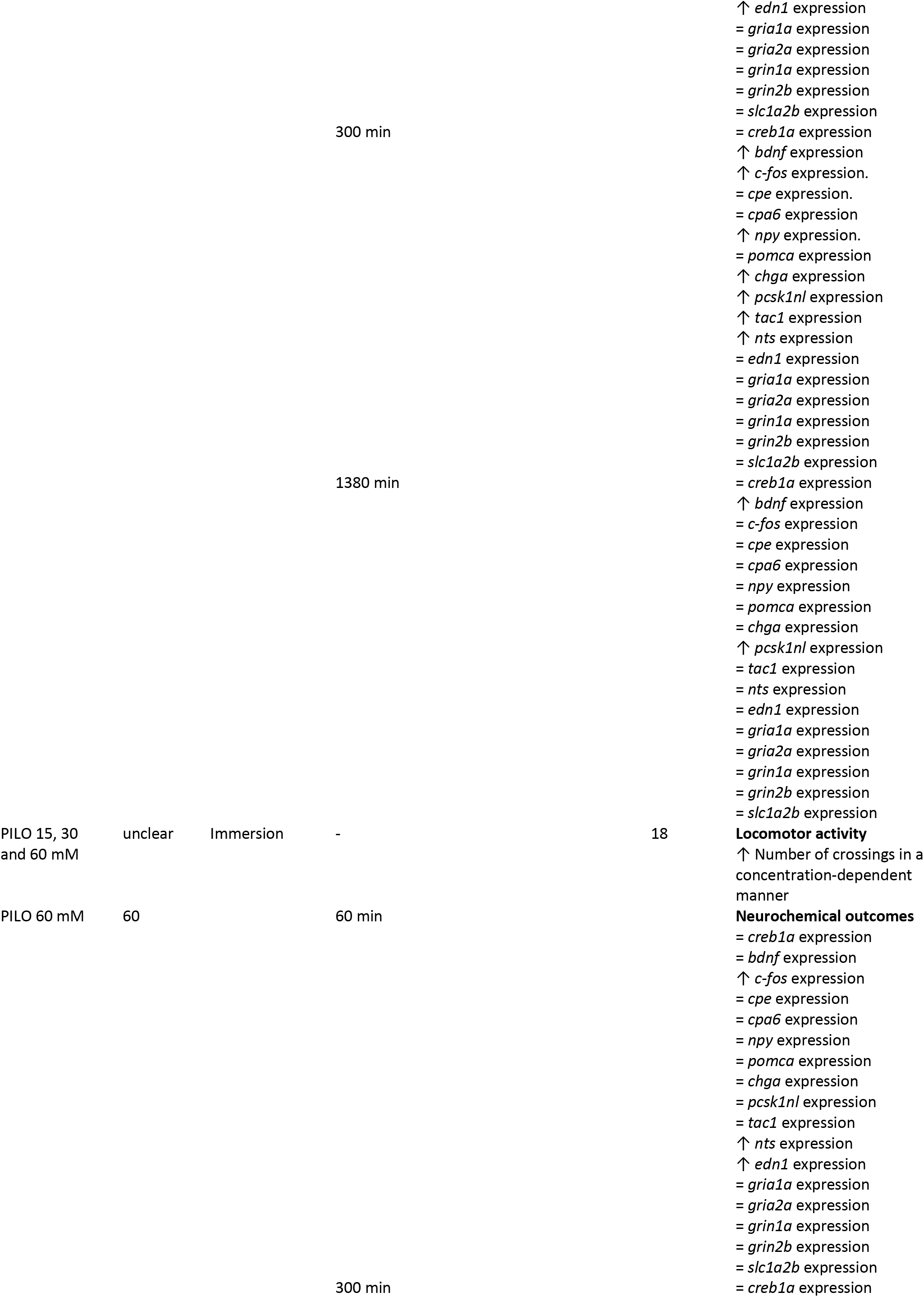
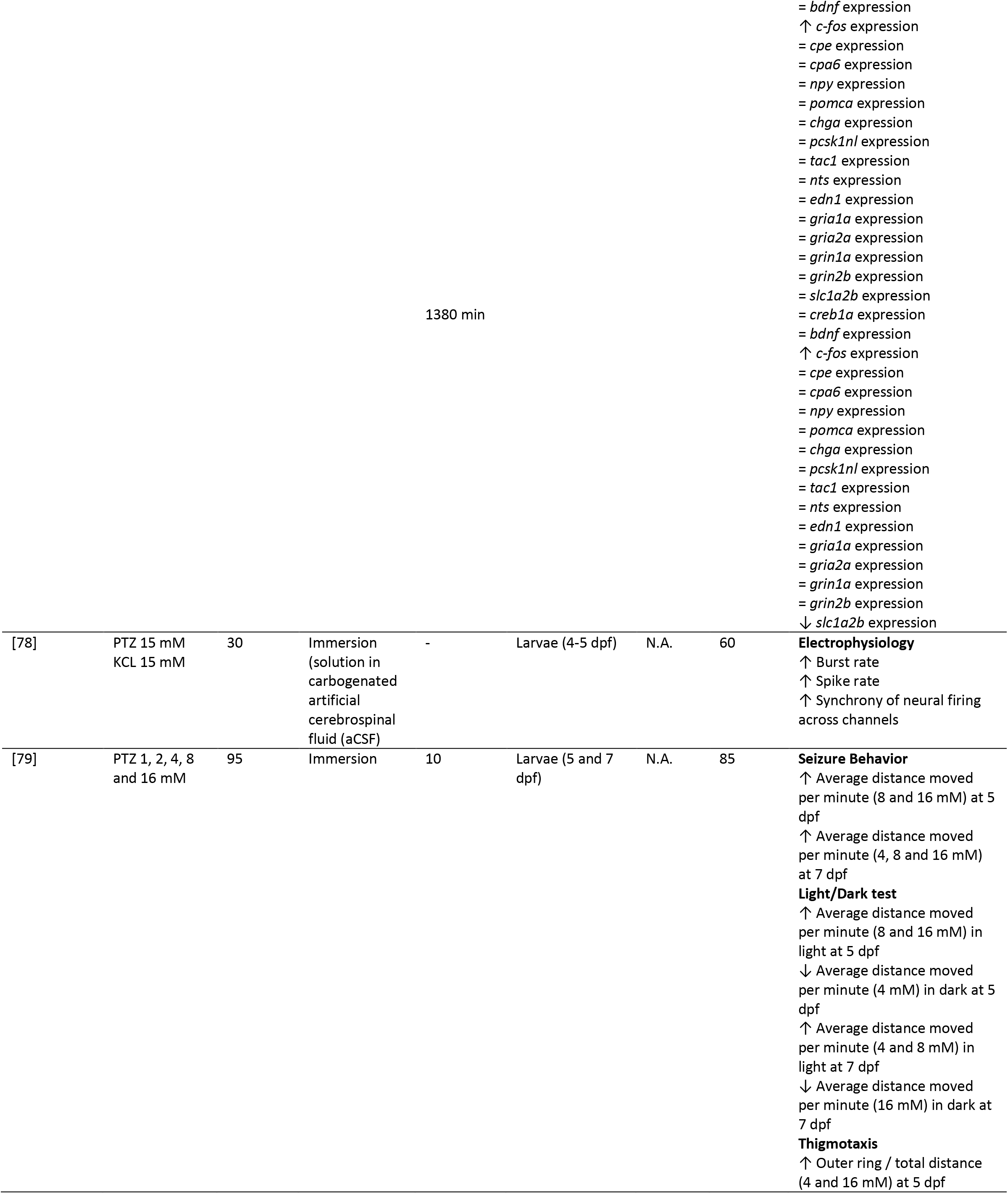
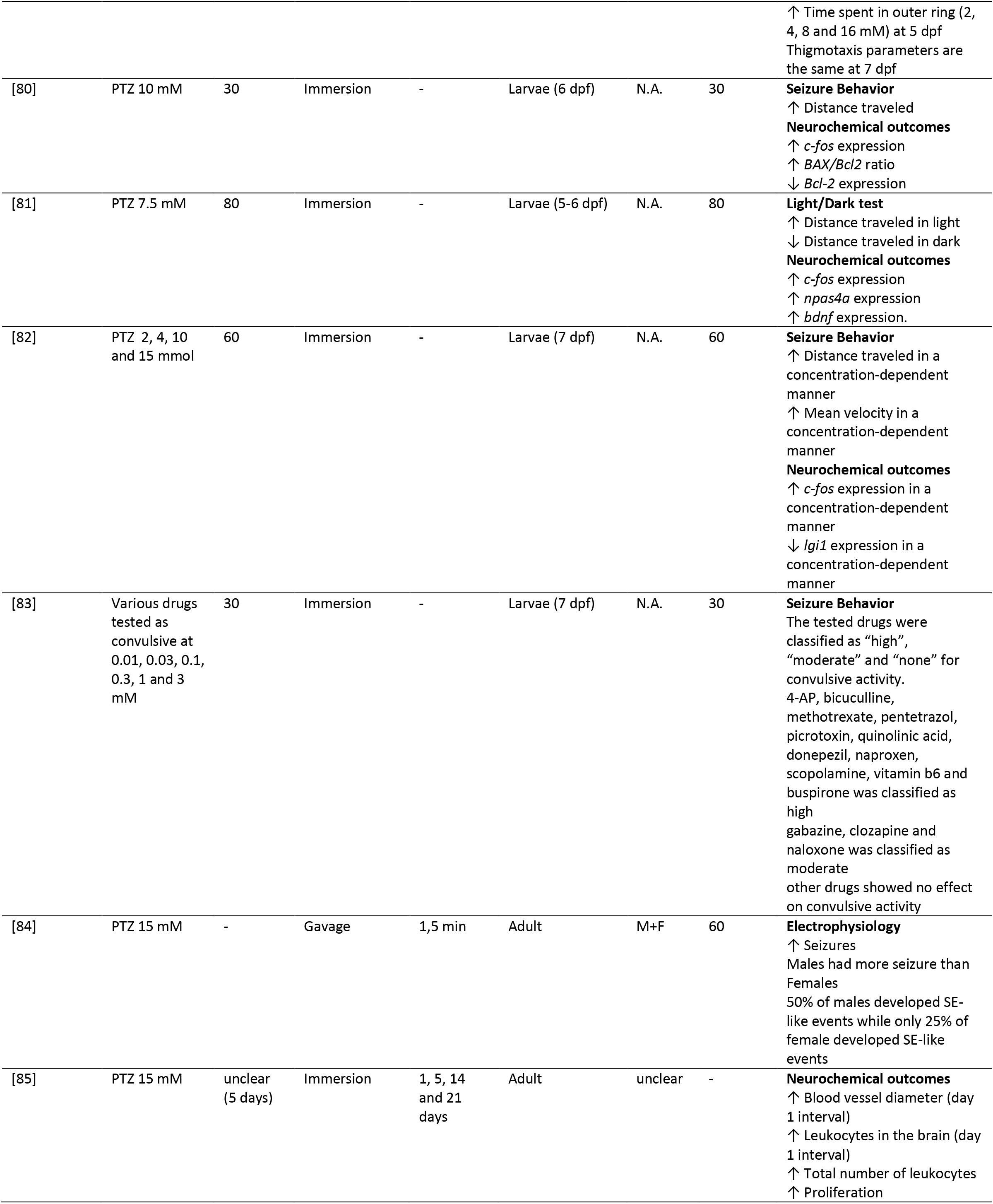
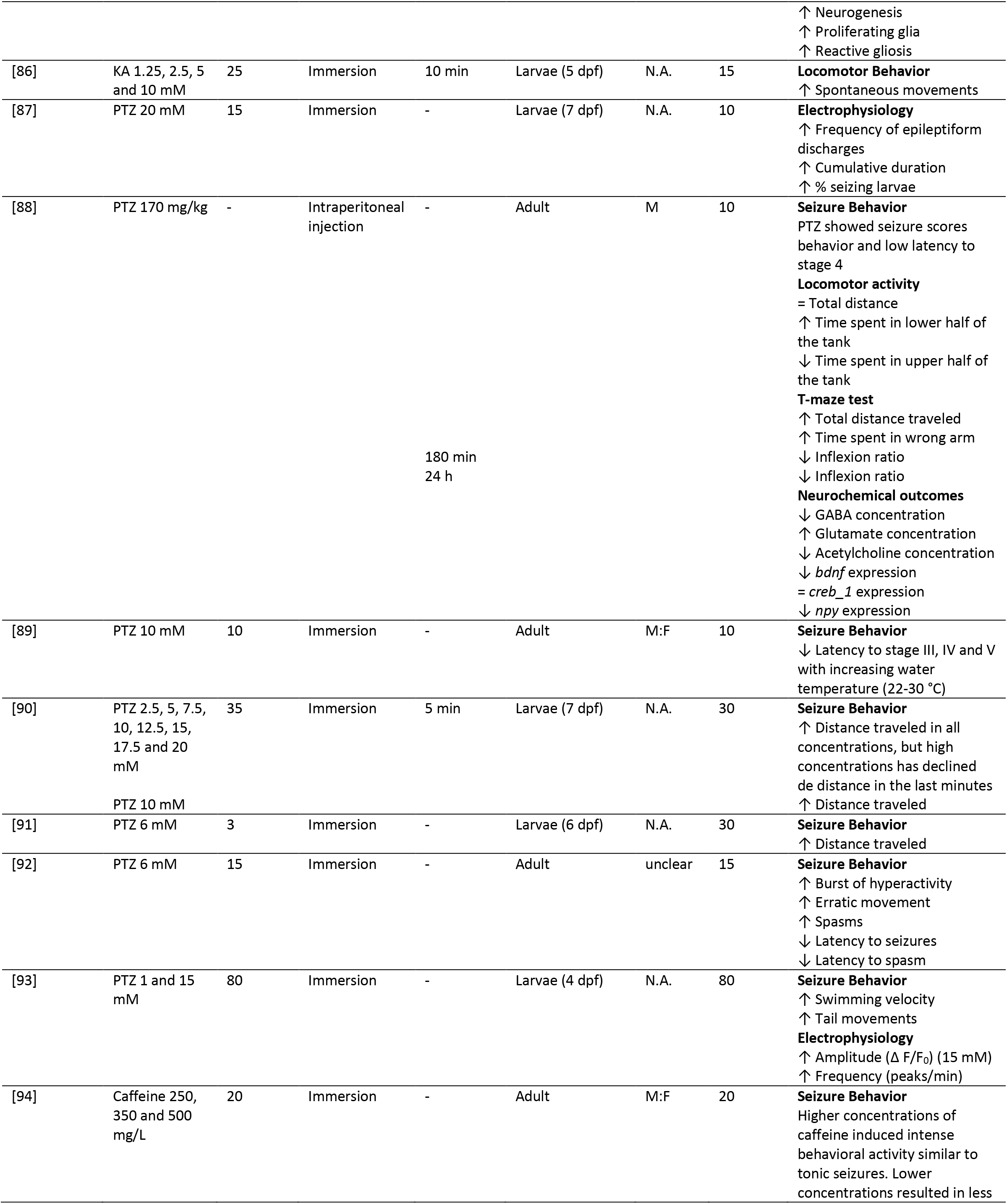
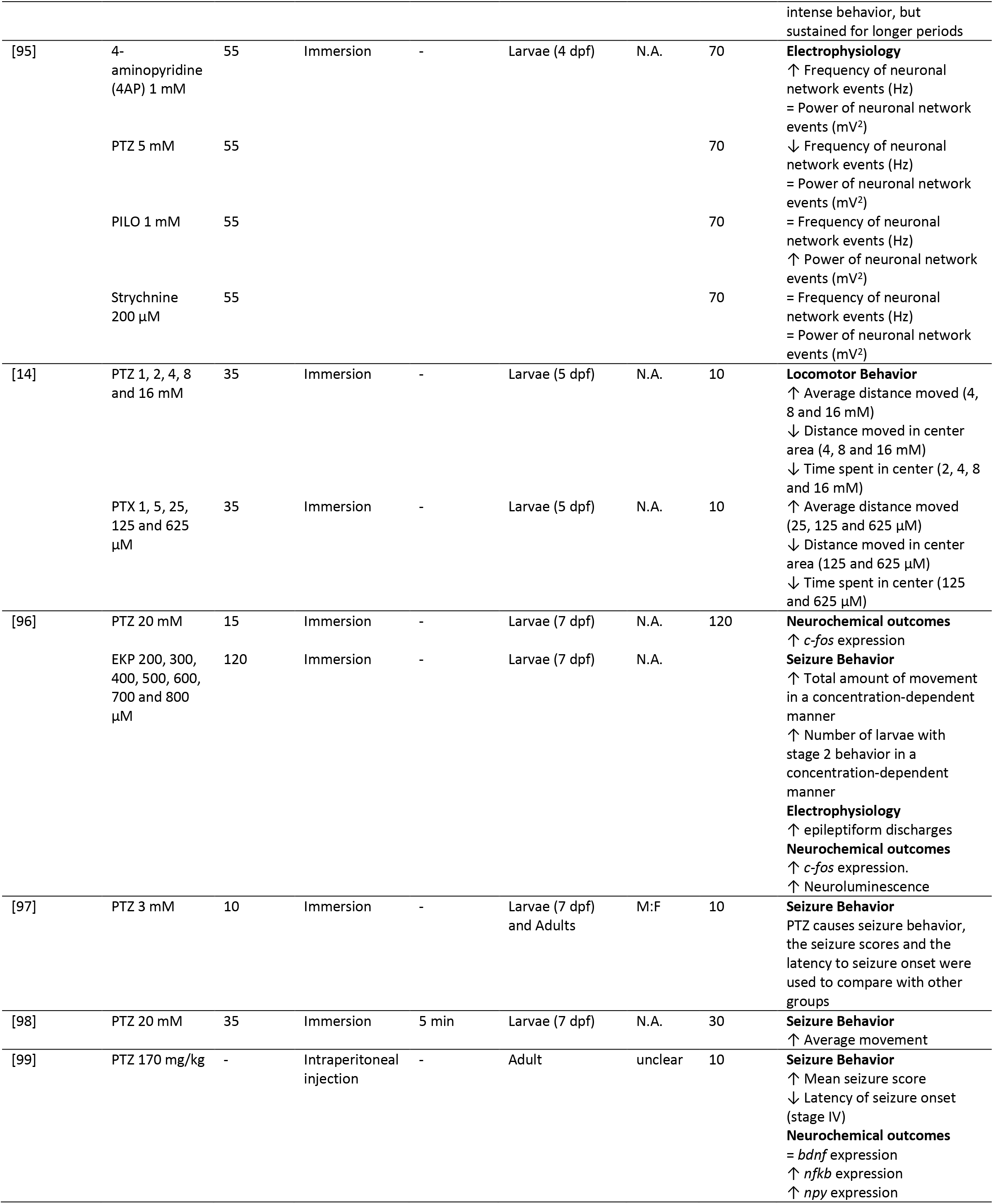
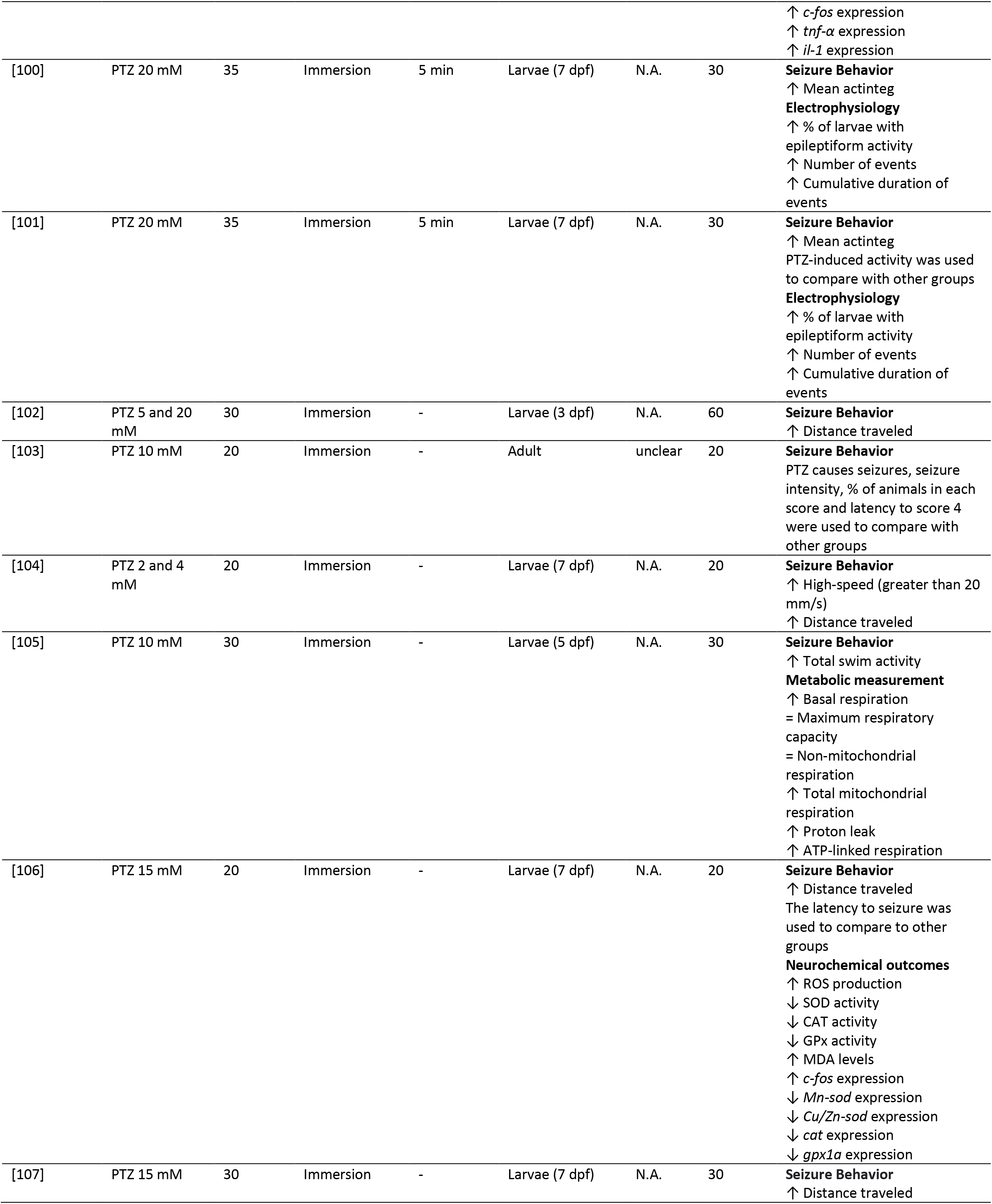
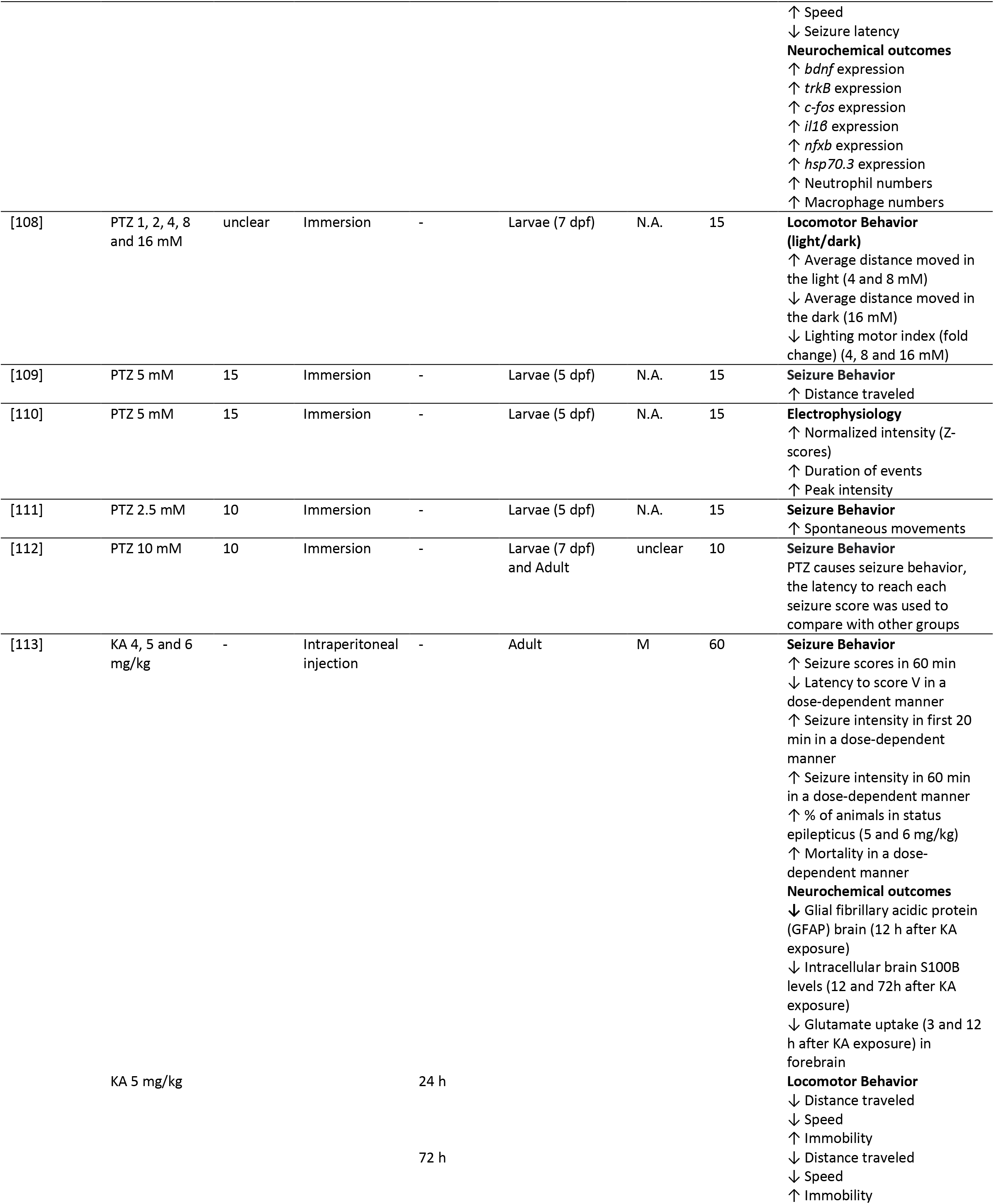
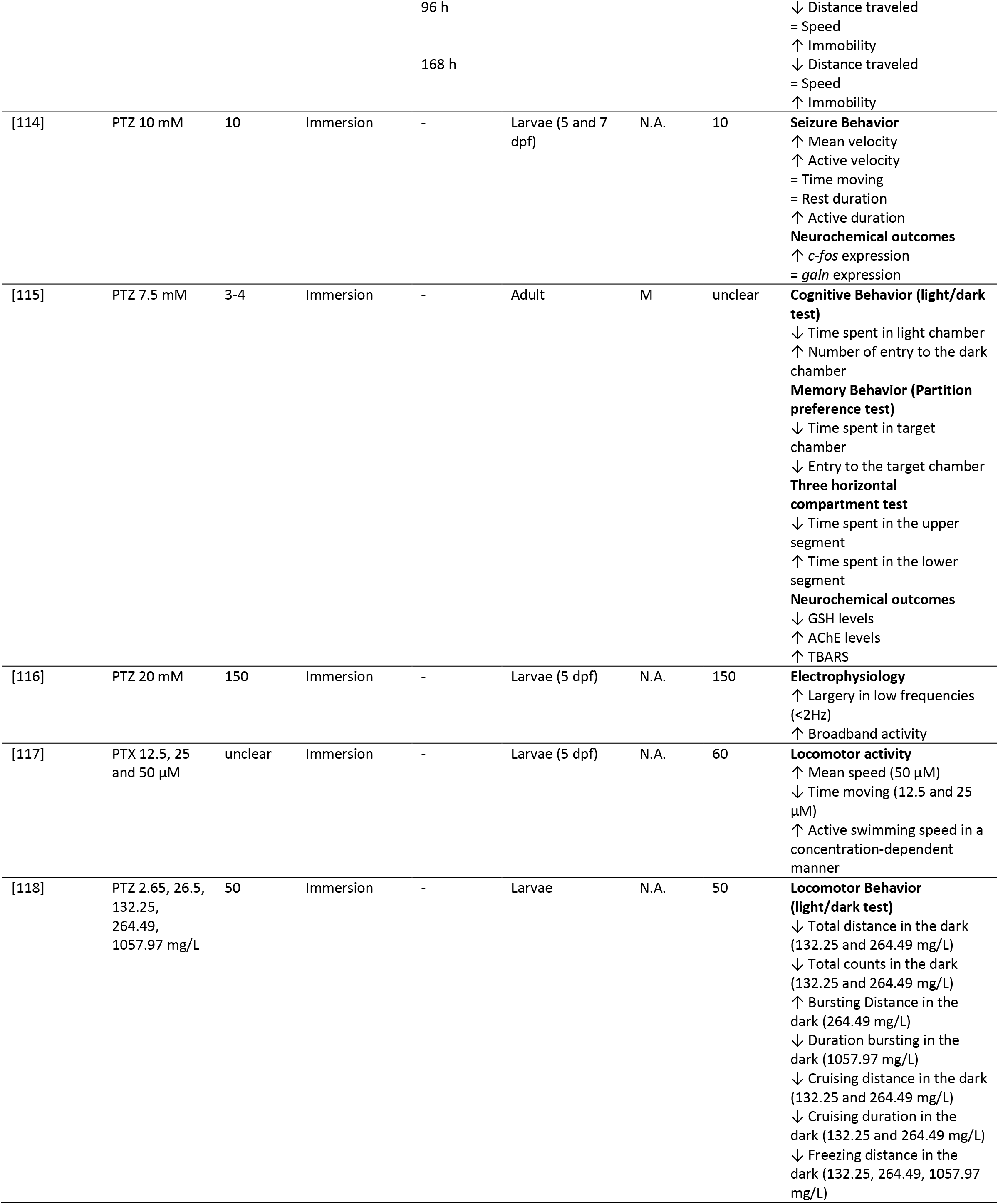
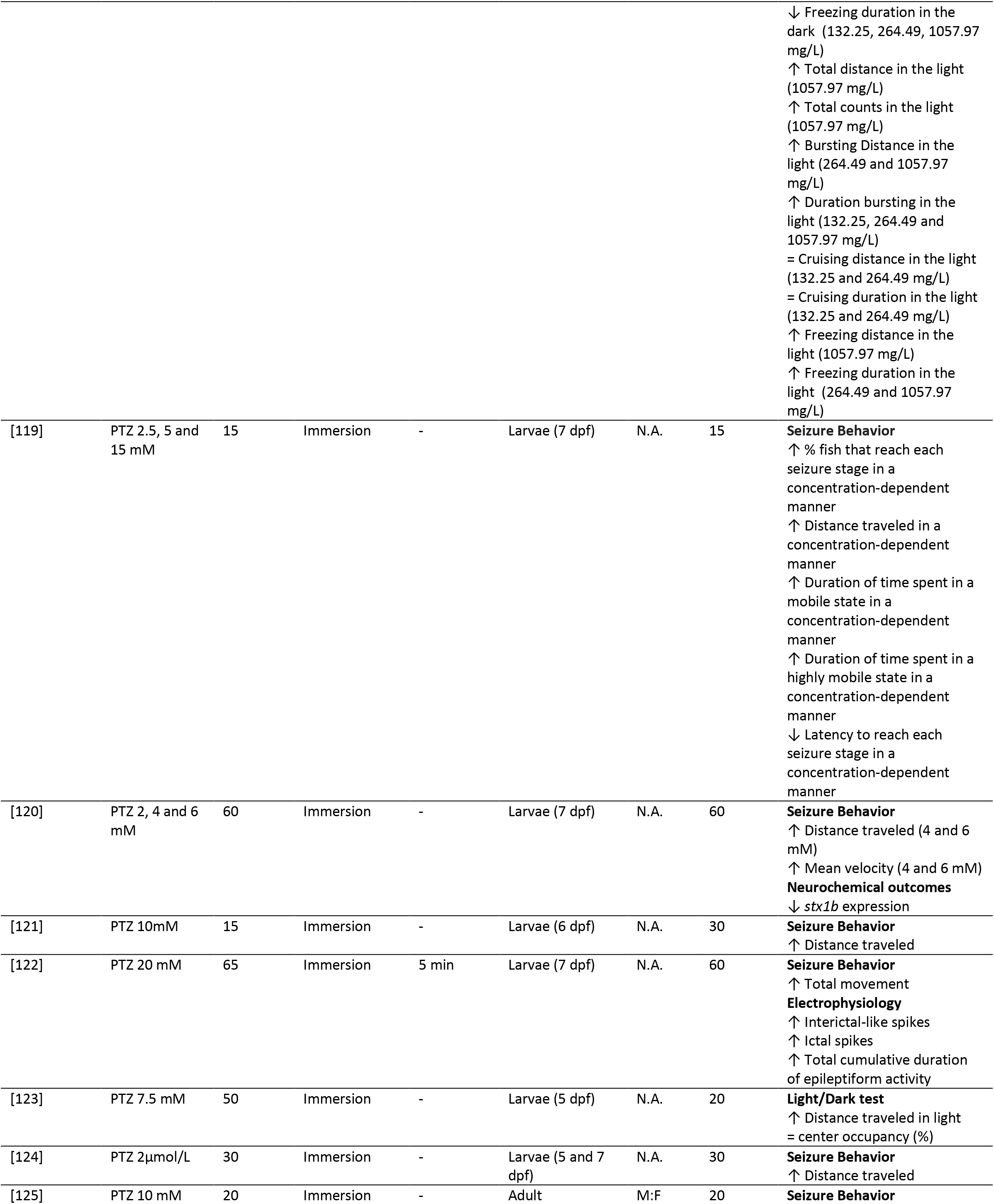
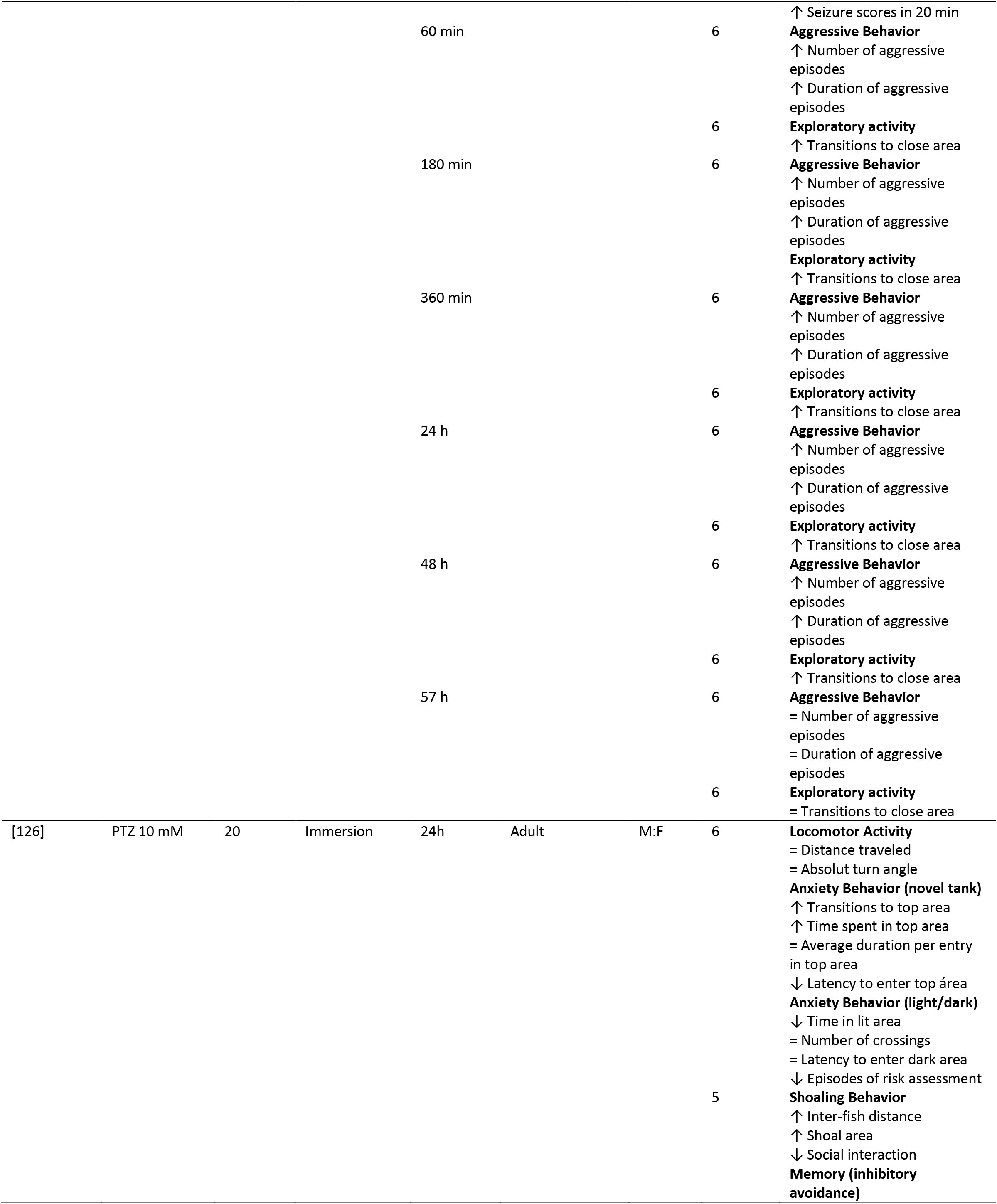
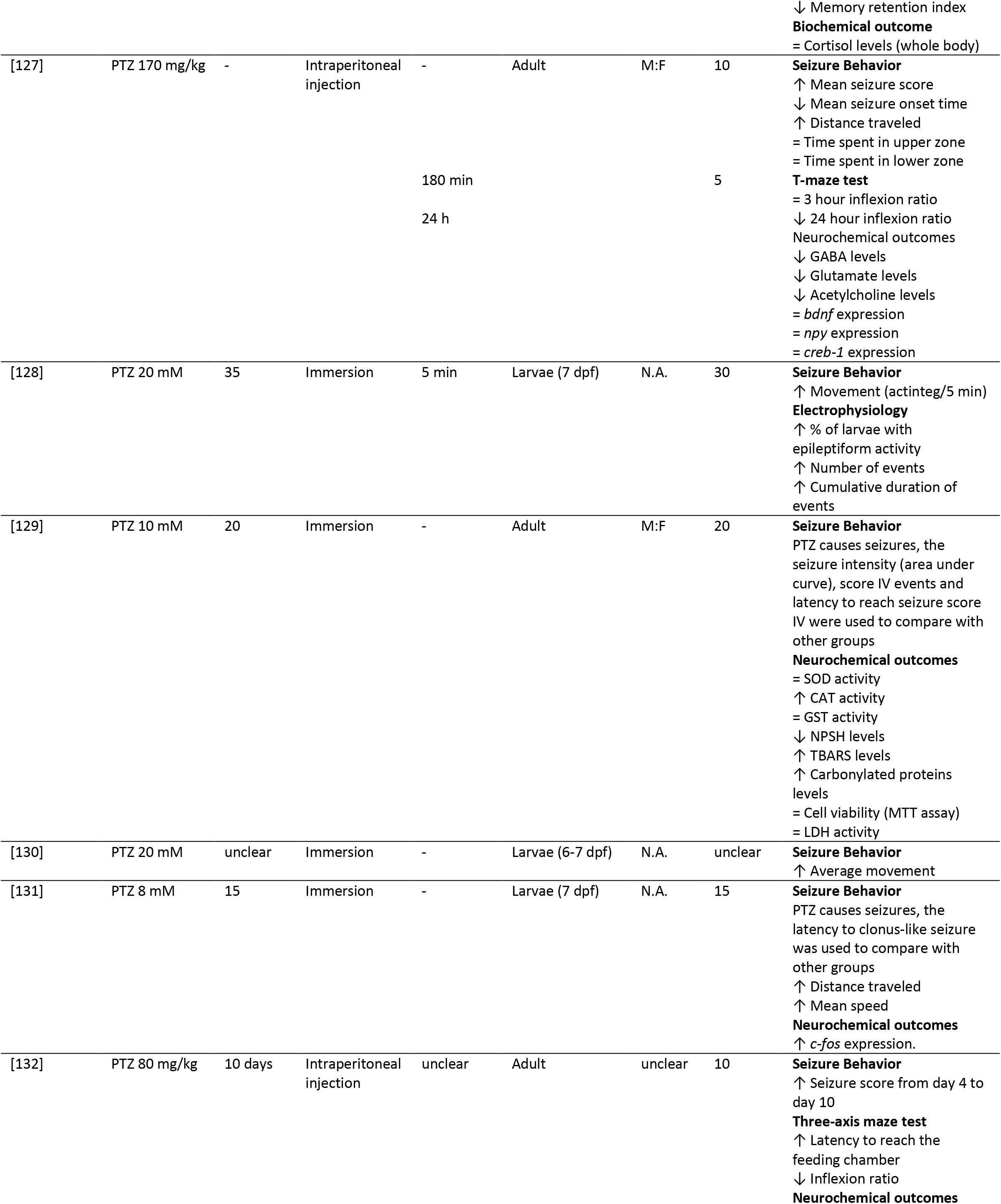
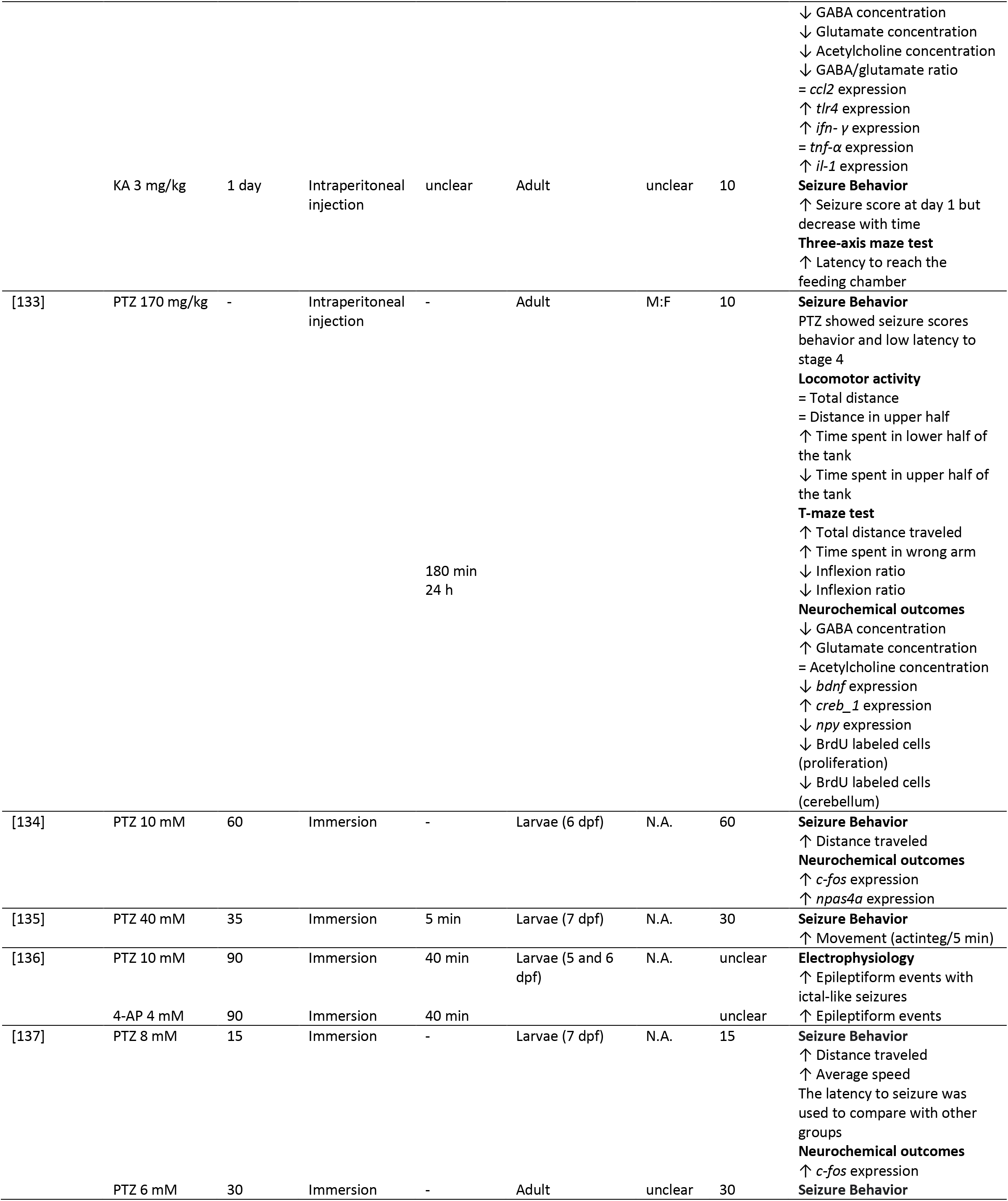
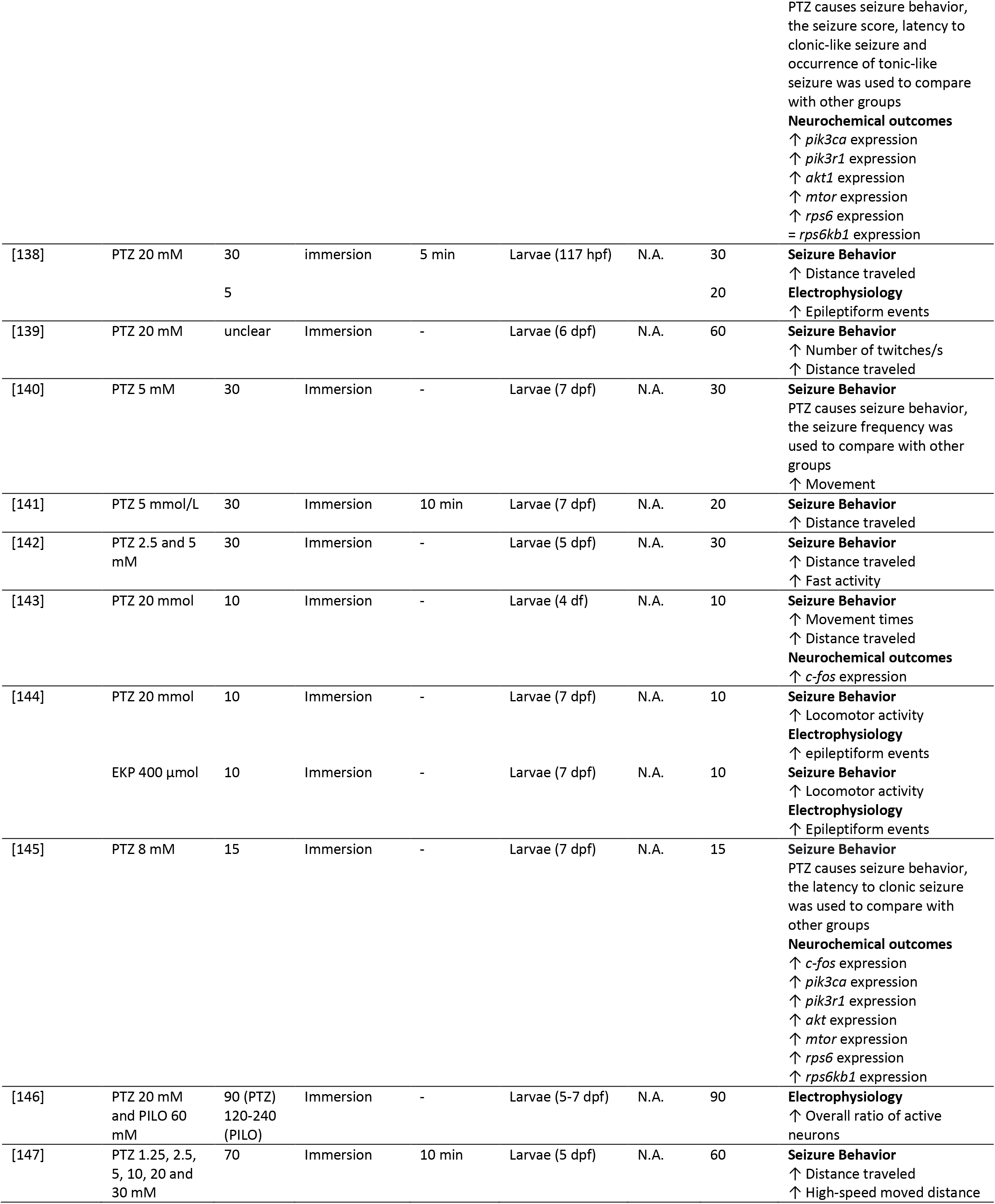
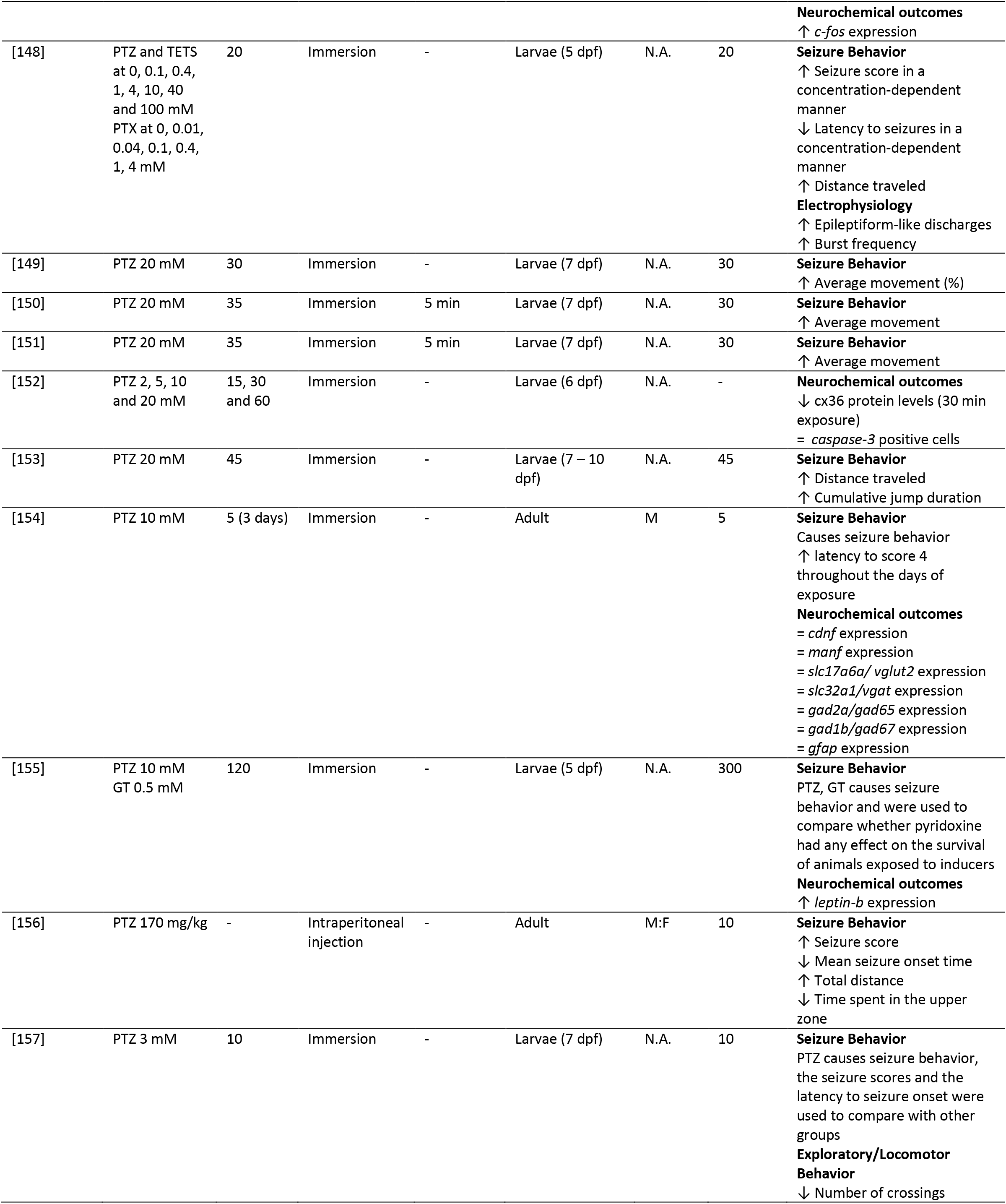
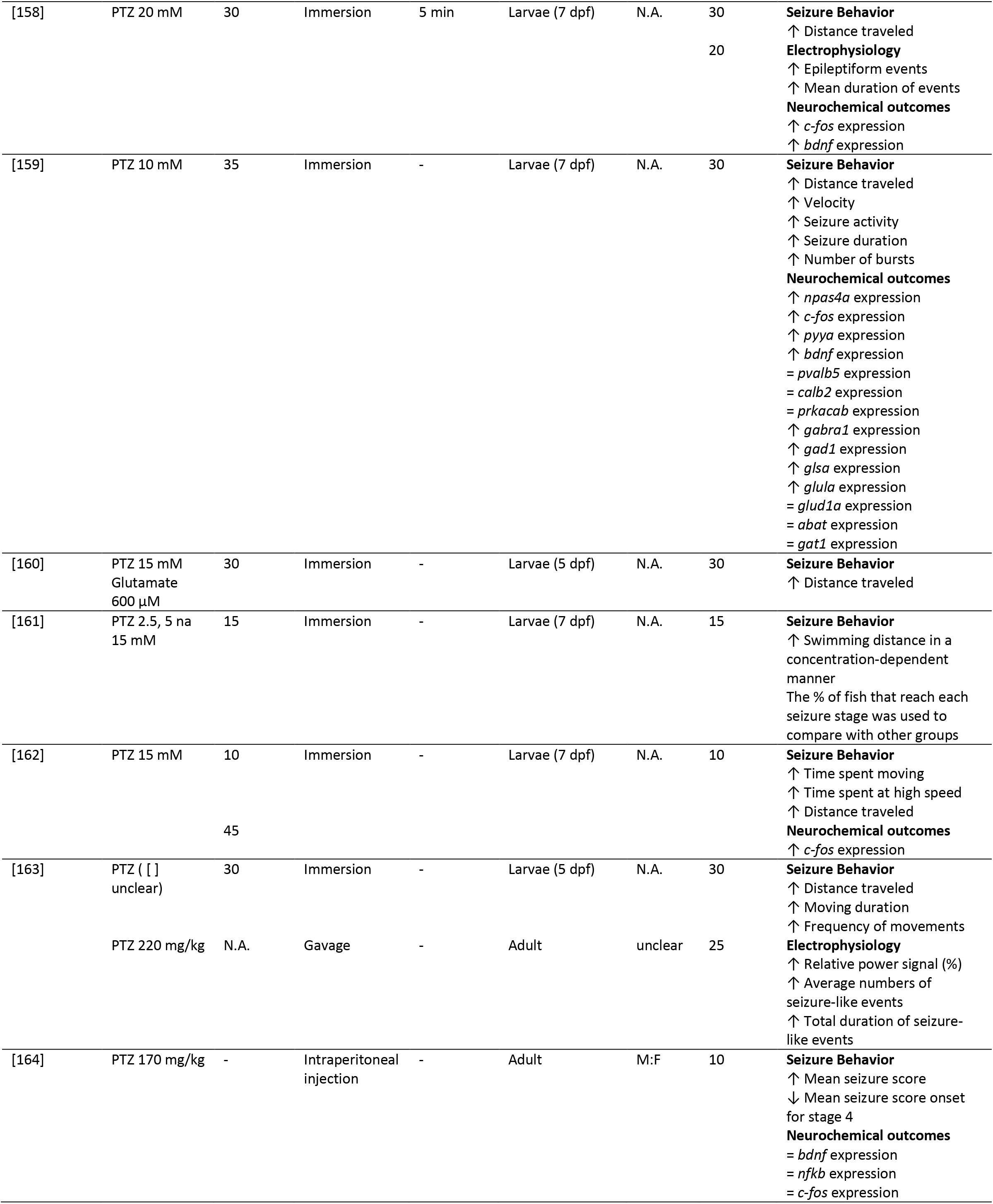
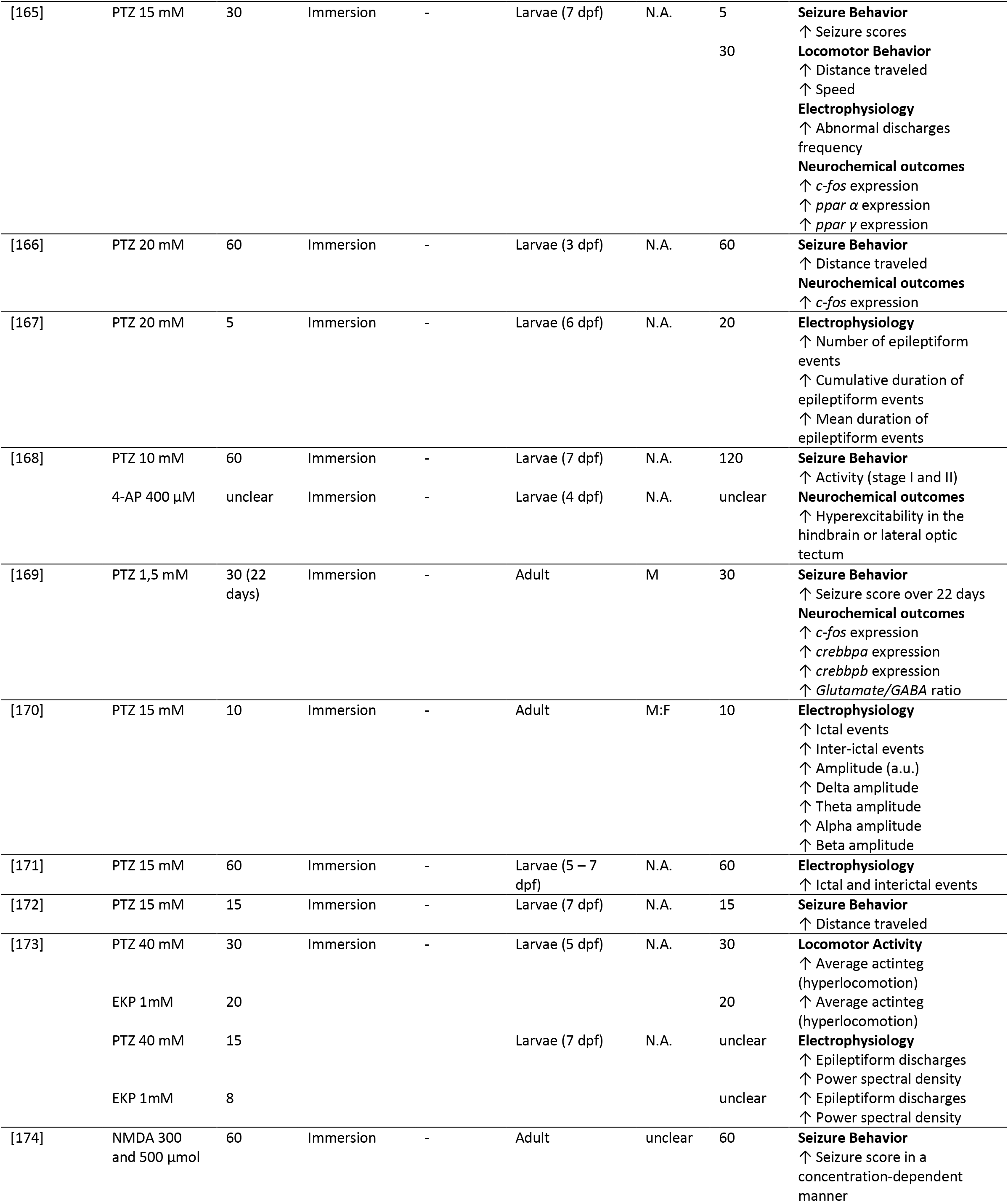
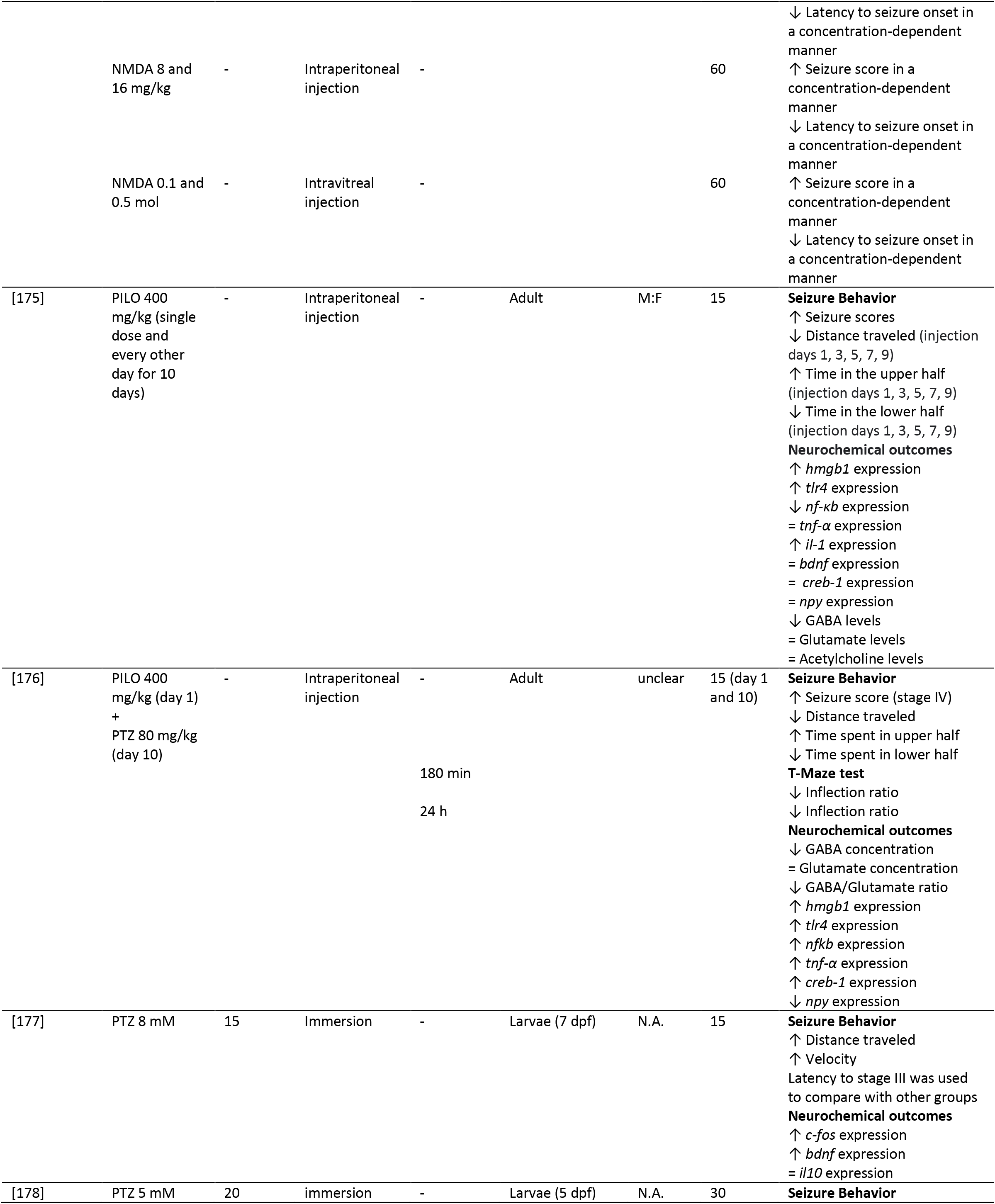
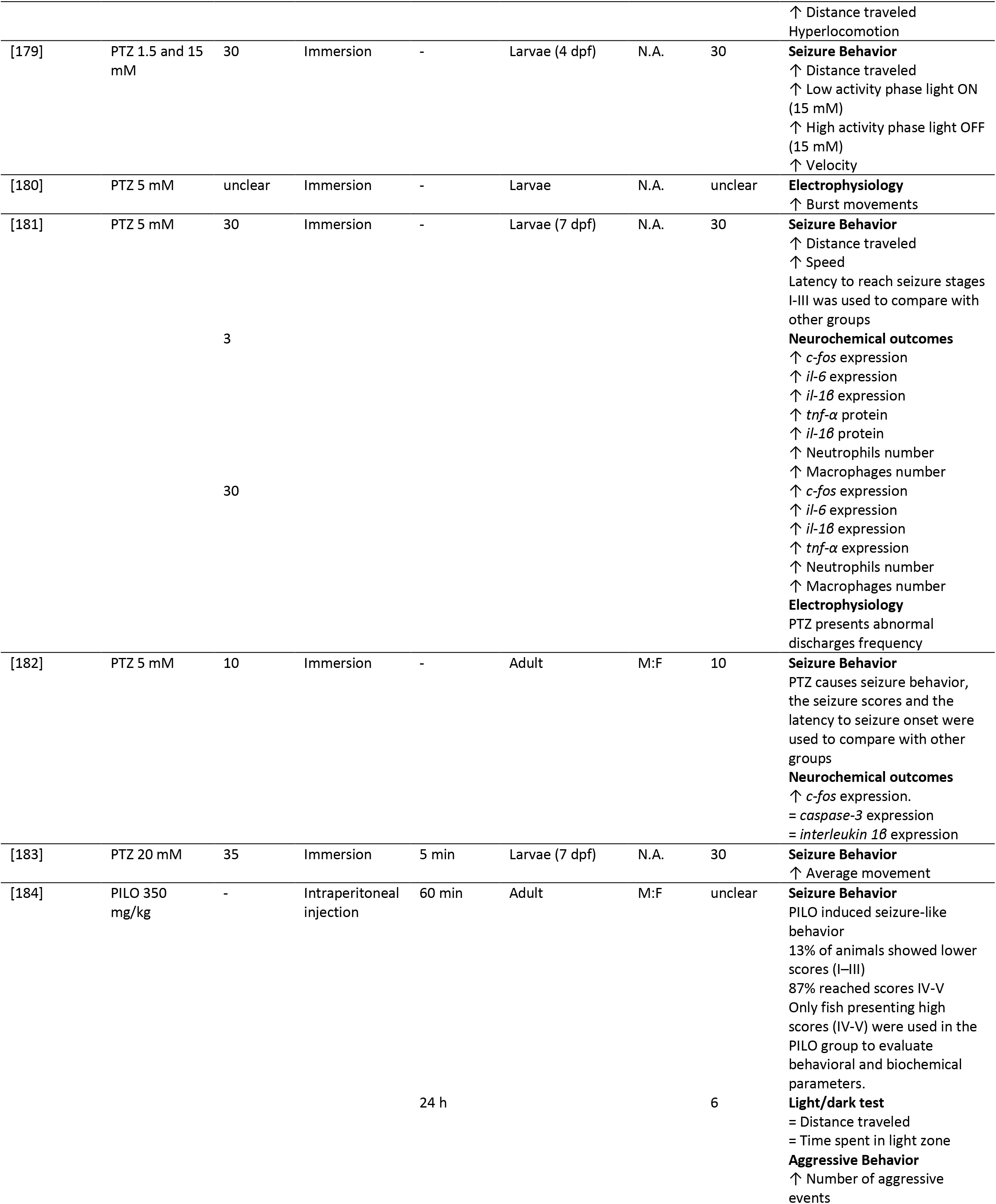
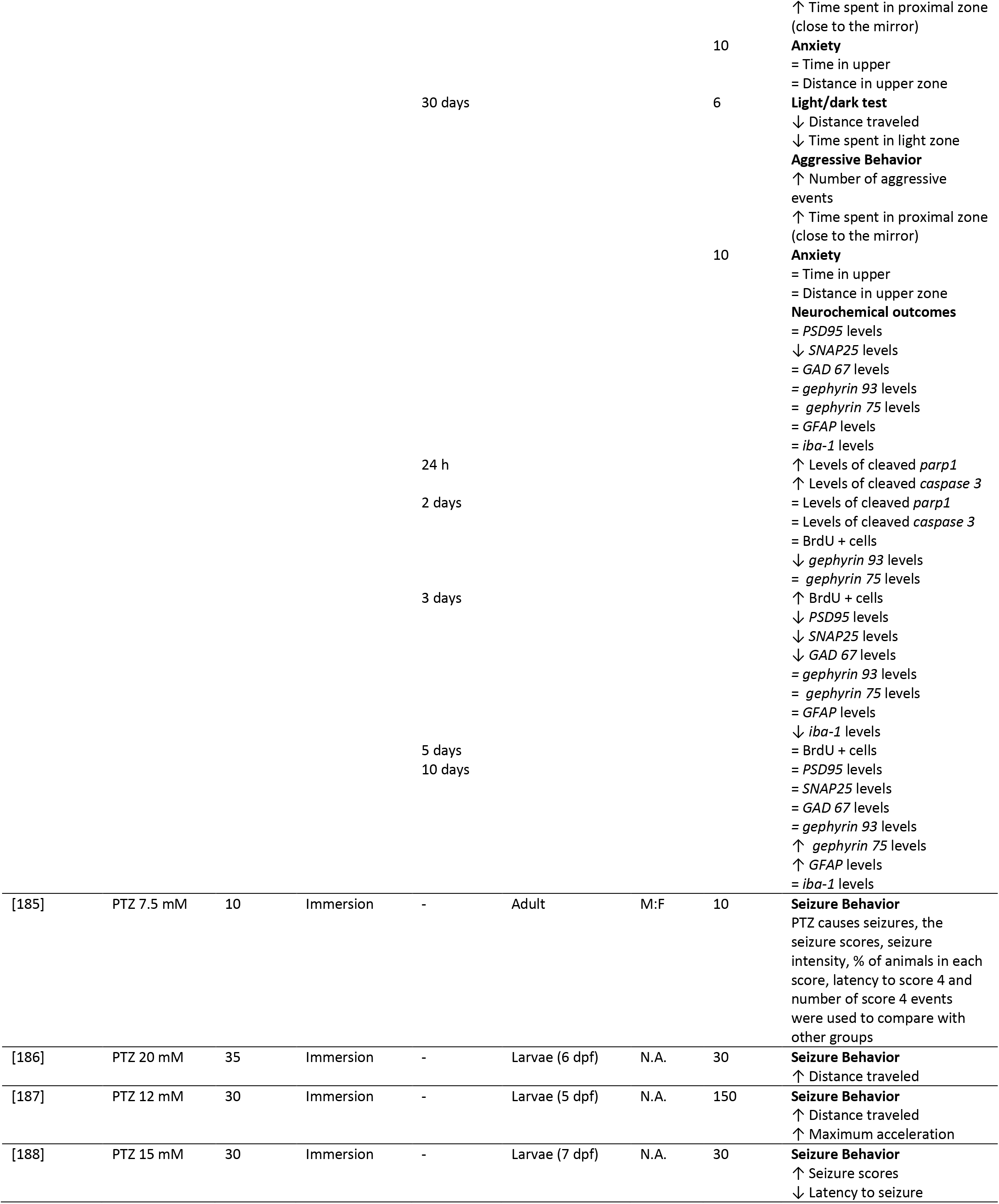
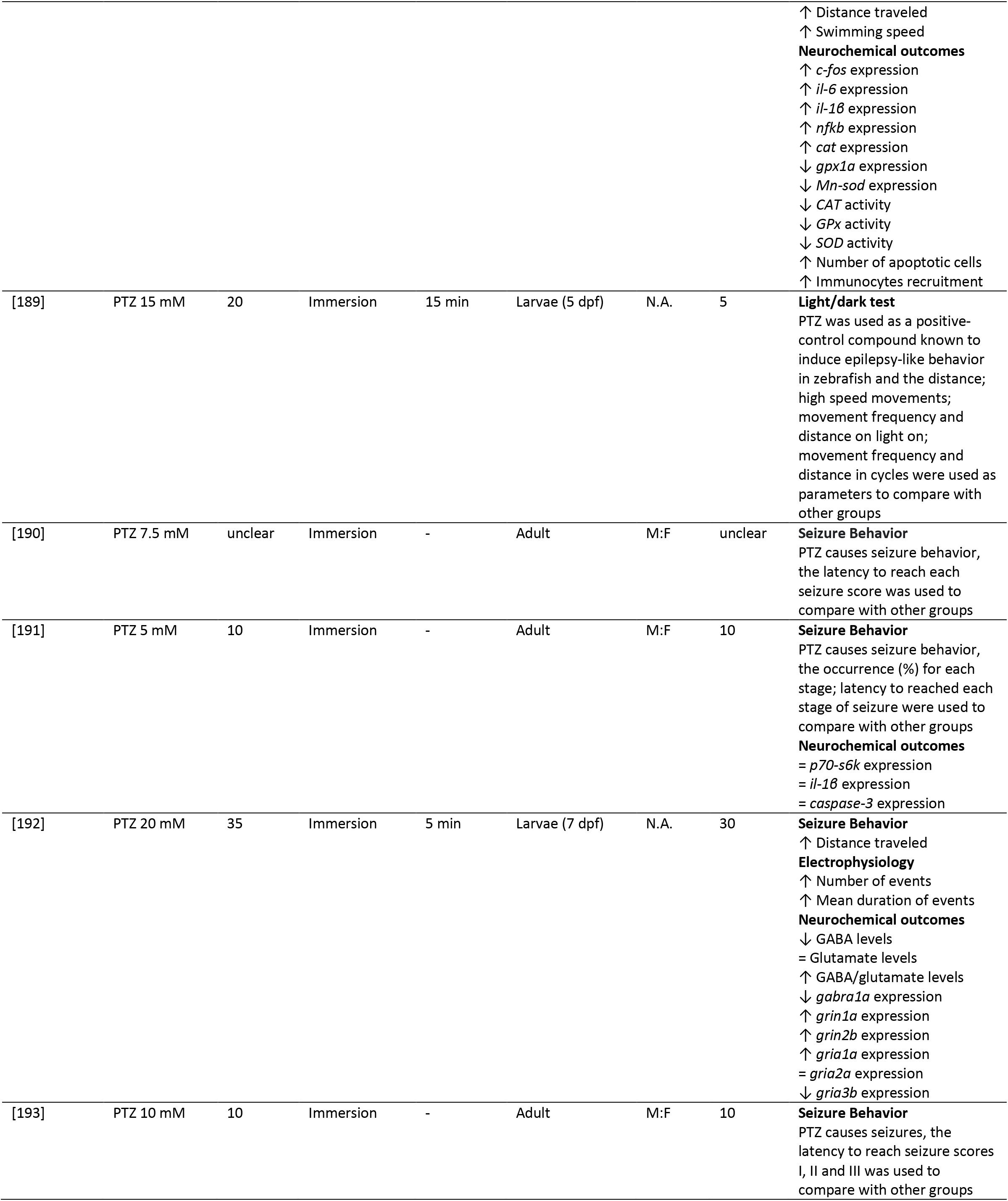
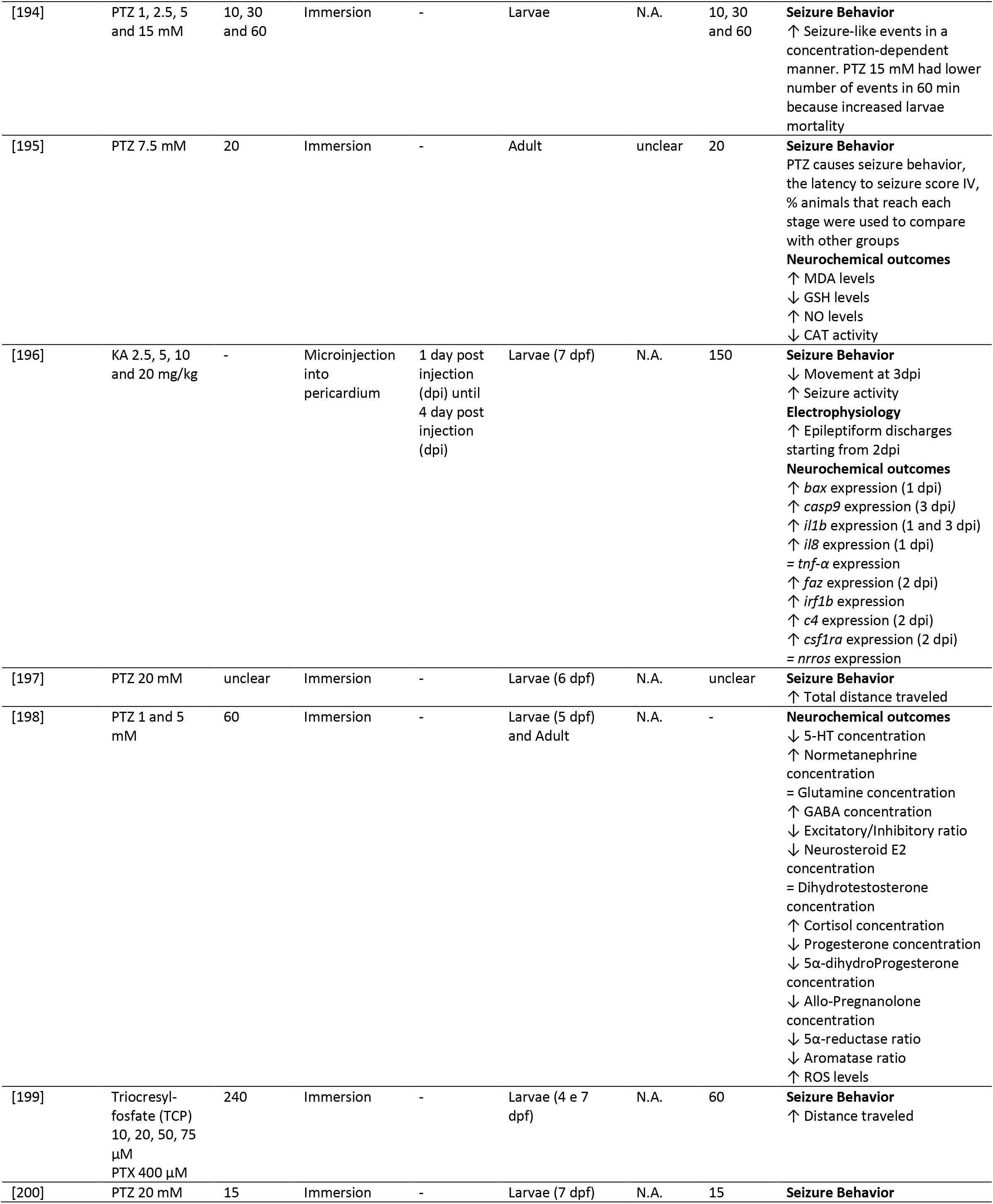
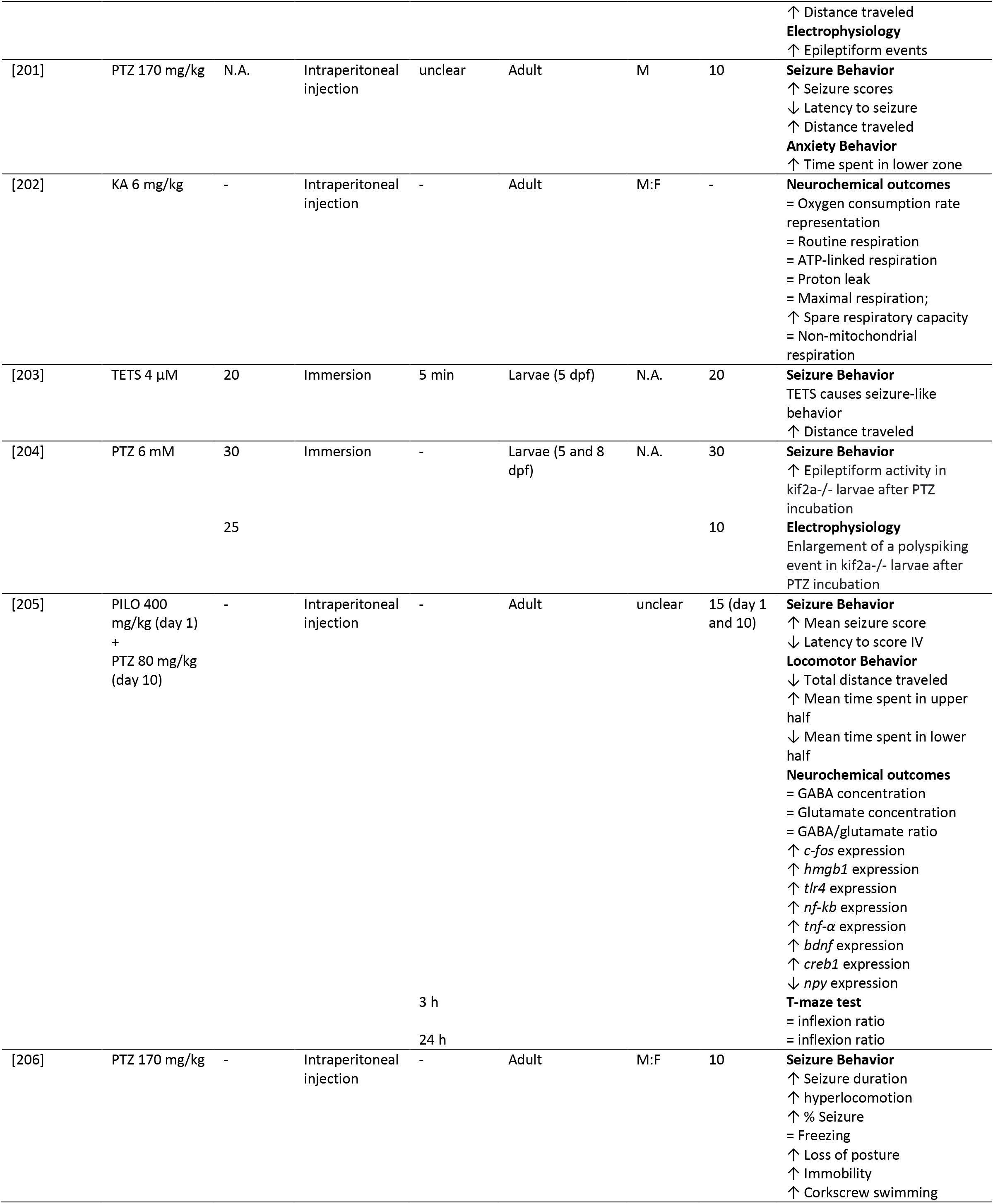
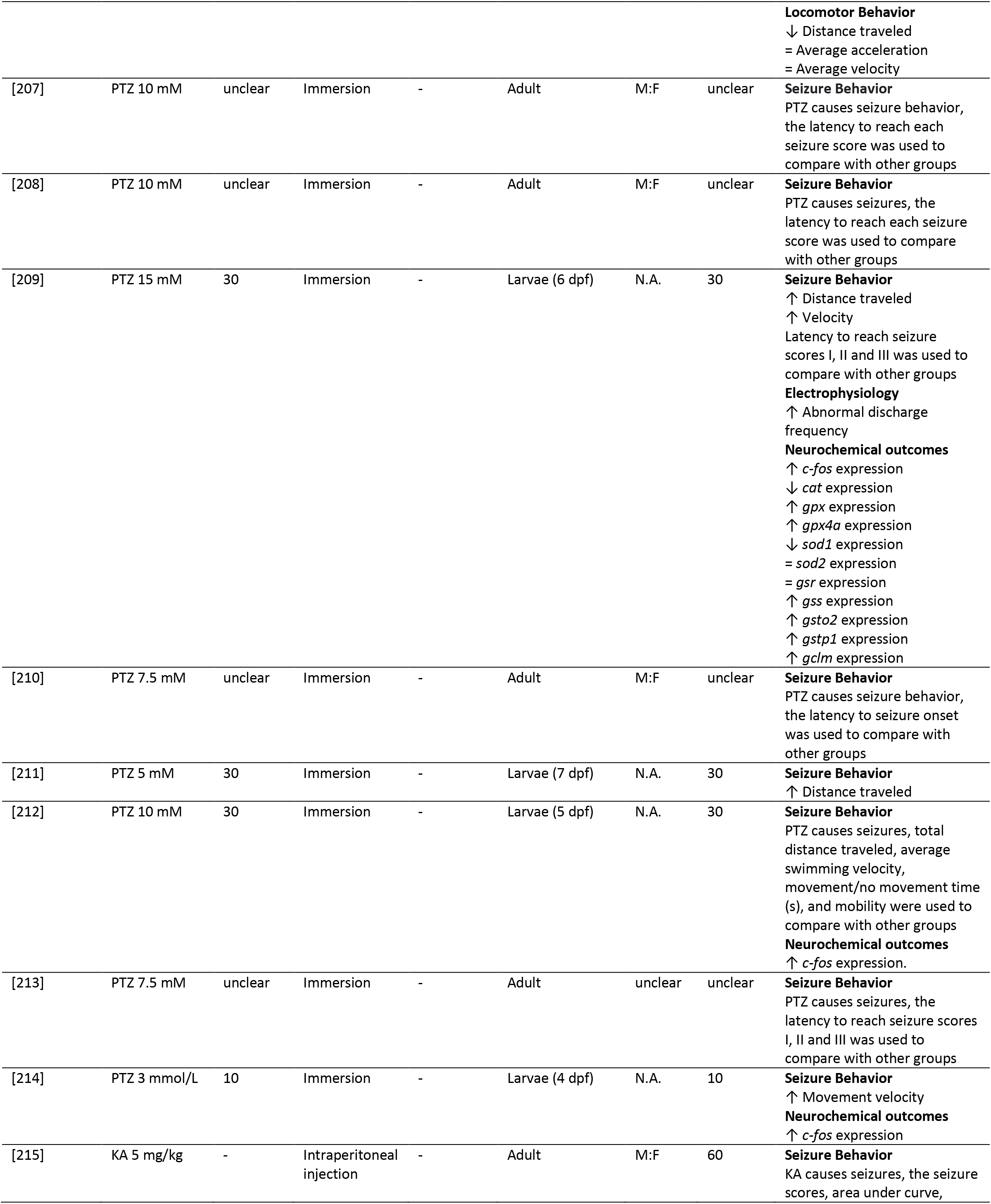
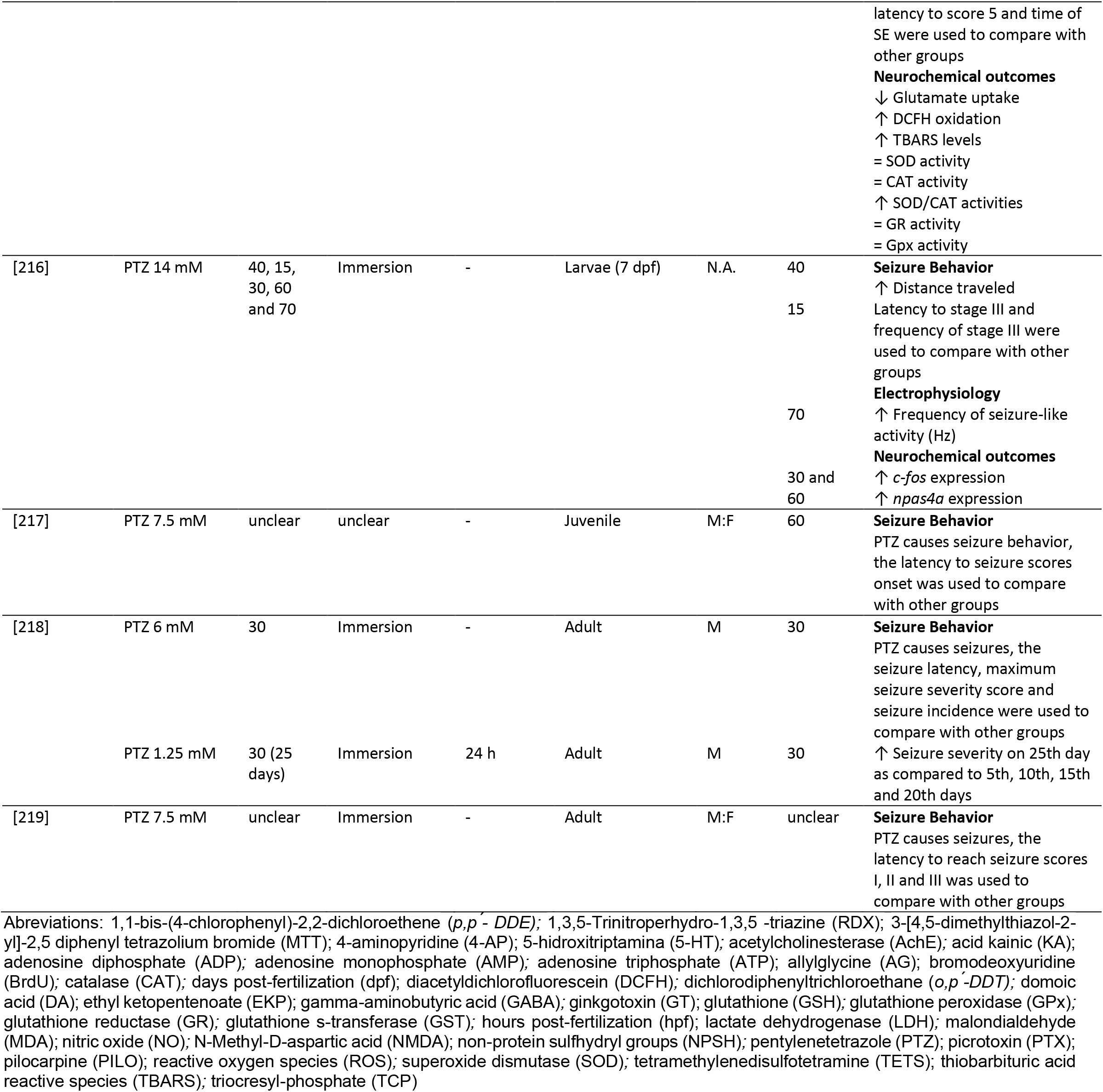
Qualitative description of studies reporting chemically-induced seizures protocols in research with zebrafish. The sex of the animals used was computed as: M, for male; F, for females; M:F, when male and female were included but tested and analyzed as a mixed group; not applicable (N.A.) for larvae and unclear when the sex of the animals was not reported. Unclear was used to address missing information in the articles. The main findings were described as: ↑, higher than the control group; ↓, lower than the control group; =, no difference when compared to the control group.

The route of administration of epileptic seizure inducers in the selected articles was mostly by immersion in a solution containing the drug (n = 178, 88.56%) or intraperitoneal injection (n = 23, 11.44%). PTZ was mainly administered by immersion, with 166 studies out of 180 that used this seizure-inducer employing this route of administration, which corresponds to 92.22%. On the other hand, most exposures to kainic acid used intraperitoneal injection as the route of administration (n = 7, 63.63%), 3 used immersion, and 1 used microinjection into pericardium. Another seizure inducer that is commonly administered by intraperitoneal (i.p.) injection is pilocarpine, where 4 studies (50%) used this method of exposure, while another 4 (50%) used immersion as the form of administration. As for picrotoxin, in all studies immersion was used as delivery route (n = 8, 100%). Domoic acid was administered by immersion in 3 studies, by microinjection in one, and via intraperitoneal injection in another one. For 4-aminopyridine, caffeine, and ethyl ketopentenoate, all studies used immersion as the route of administration.

As expected, the duration of exposure to seizure inducers varies between studies. In the 180 studies using PTZ, the time of exposure ranged from 2 to 390 min, with 30 min as the most frequently used exposure time (n = 37, 20.55%). Other commonly used exposure times were 10 min (n = 24, 13.33%), 15 min (n = 19, 10.55%), 60 min (n = 18, 10%), and 20 min (n = 17, 9.44%). Exposure to kainic acid via immersion ranged from 10 to 30 min in 3 studies. The duration of exposure to pilocarpine by immersion ranged between 2 and 240 min. Picrotoxin exposure duration ranged from 20 to 240 min, and in 2 studies this information was unclear.

The utilization of various doses and concentrations in chemically-induced seizure protocols are evident from the data presented in Table 1. It is noteworthy that different research groups employ a range of inducers, resulting in variations in the experimental conditions. For PTZ, for example, we found a variation between less than 1 and more than 60 mM, and the most commonly used concentration in cases where immersion was the chosen route of administration was 20 mM (n = 40, 22.22%), followed by 10 mM (n = 39, 21.66%) and 15 mM (n = 35, 19.44%). For studies that used intraperitoneal injection, the most commonly used dose was 170 mg/kg (n = 8, 4.44%). Other doses used were 220 and 225 mg/kg, in addition to the dose of 80 mg/kg administered for 10 days. In the case of kainic acid, intraperitoneal injection doses ranged from 5 or 6 mg/kg (n = 3, 27.27% for 5 mg/kg and n = 5, 45.45% for 6 mg/kg). Pilocarpine was used mainly at concentrations of 15, 30 and 60 mM when administered by immersion, with 2 studies for each concentration, and the most common intraperitoneal injection dose was 400 mg/kg. To see all the concentrations of all seizure inducers, please check Table 1.

Most of the available studies were carried out with zebrafish in the early developmental stages, with 136 studies on embryos/larvae, corresponding to 67.66%, against 67 studies on adults (33.33%). Only 3 studies used zebrafish in the juvenile stage (1.49%). In larvae, most studies were conducted at 7 days post-fertilization (dpf) (n = 80, 58.82%), followed by 5 dpf (n = 34, 25%) and 6 dpf (n = 19, 13.97%) (Table 1).

The vast majority of studies on epilepsy using zebrafish as a model organism are studies of acute exposure to a chemoconvulsant to assess epileptic seizures for the discovery of new compounds through behavioral assessment. 178 studies (88.55%) describe behavioral analyses, while 81 (40.29%) showed neurochemical outcomes such as gene expression and oxidative stress status, and 44 (21.89%) reported electrophysiological results. Among the behavioral outcomes most commonly assessed during epileptic seizures are distance traveled (n=89, 50%), latency to reach each seizure score (n = 60, 33.71%) and ratings of epileptic seizure scores (n = 42, 23.59%). In addition, 44 studies use electrophysiology to verify occurrence, frequency and duration of epileptiform discharges.

Within neurochemical analyses, changes in genes involved in epileptogenesis and neuronal activity are widely evaluated. The most evaluated gene among the studies is *c-fos* (n = 44, 54.32%), followed by *bdnf* (n = 15, 18.51%). Other genes appear in fewer studies, such as *npy, tnf-α, creb_1, tlr4, nfkb* and *caspase-3*. Additionally, glutamate and gamma-aminobutyric acid (GABA) levels are also seen in 11 and 10 studies, respectively. Oxidative stress appears in 8 studies, with analyses such as levels of lipid peroxidation, and activity or expression of enzymes such as superoxide dismutase (SOD) and catalase (CAT).

Co-authorship network analysis identified 76 clusters of researchers that use chemically-induced seizure protocols in their labs across the globe based on the studies included in this review (Fig. 2). An interactive version of the co-authorship network is available at https://tinyurl.com/2g9kgyu6.

**Fig. 2.**
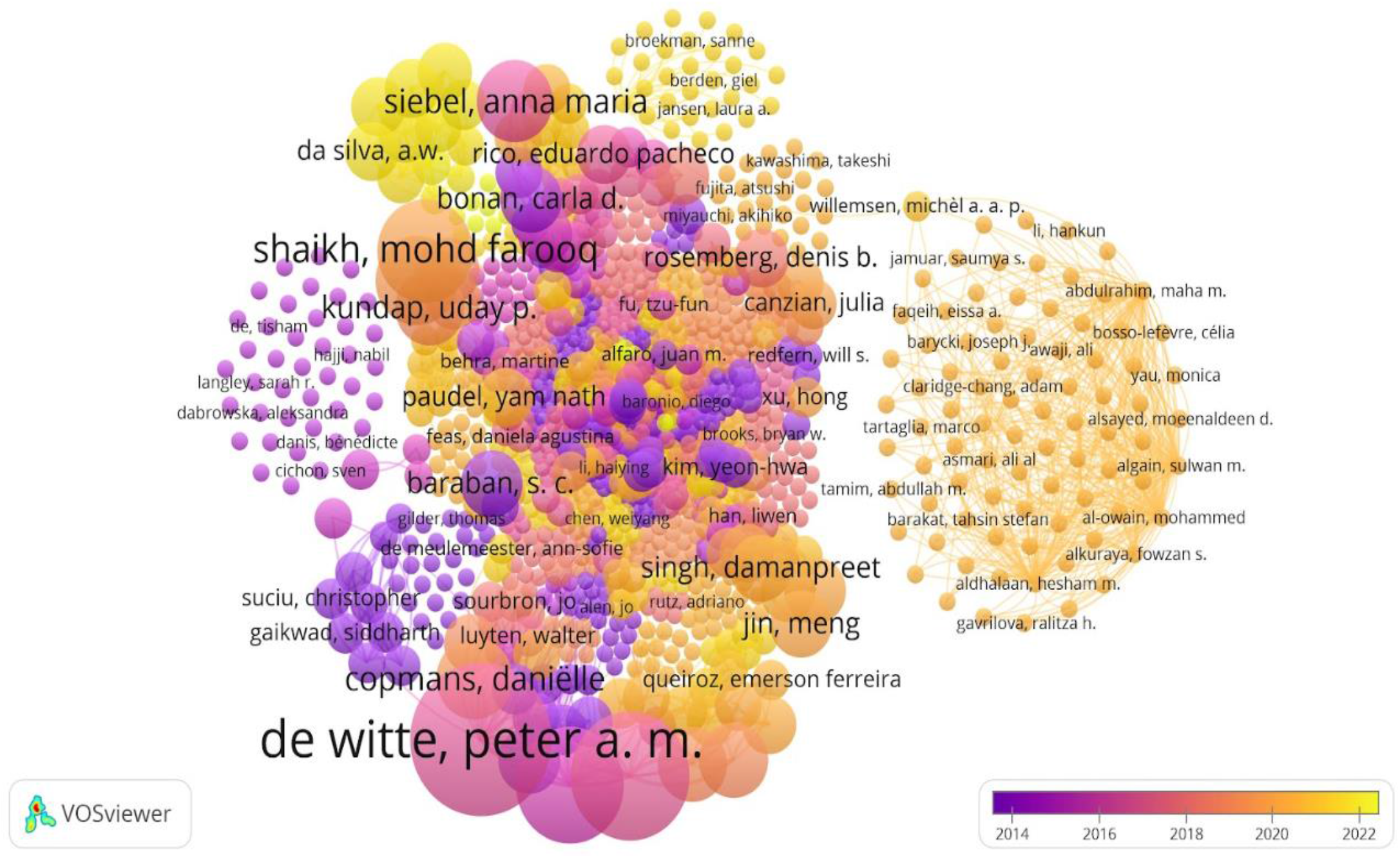
Co-authorship network analysis of researchers that authored studies implementing chemically-induced seizures in zebrafish. Authors are color-coded from violet (older studies) to yellow (more recent studies), indicating the average publication year of the studies published by each researcher. The size of the circles represents the number of studies published by each author. The distance between the two circles indicates the correlations between researchers.

### Risk of bias and reporting quality

A sample of 100 studies (49.75%) was randomly chosen for risk of bias and reporting quality assessment (figure 3). About 94% of the studies were rated as presenting a low risk of bias for selective reporting, 67% for baseline characteristics of the animals, and 54% for blinding. Randomization procedures and incomplete data were rated as unclear in 81% and 68% of the studies, respectively.

**Fig. 3.**
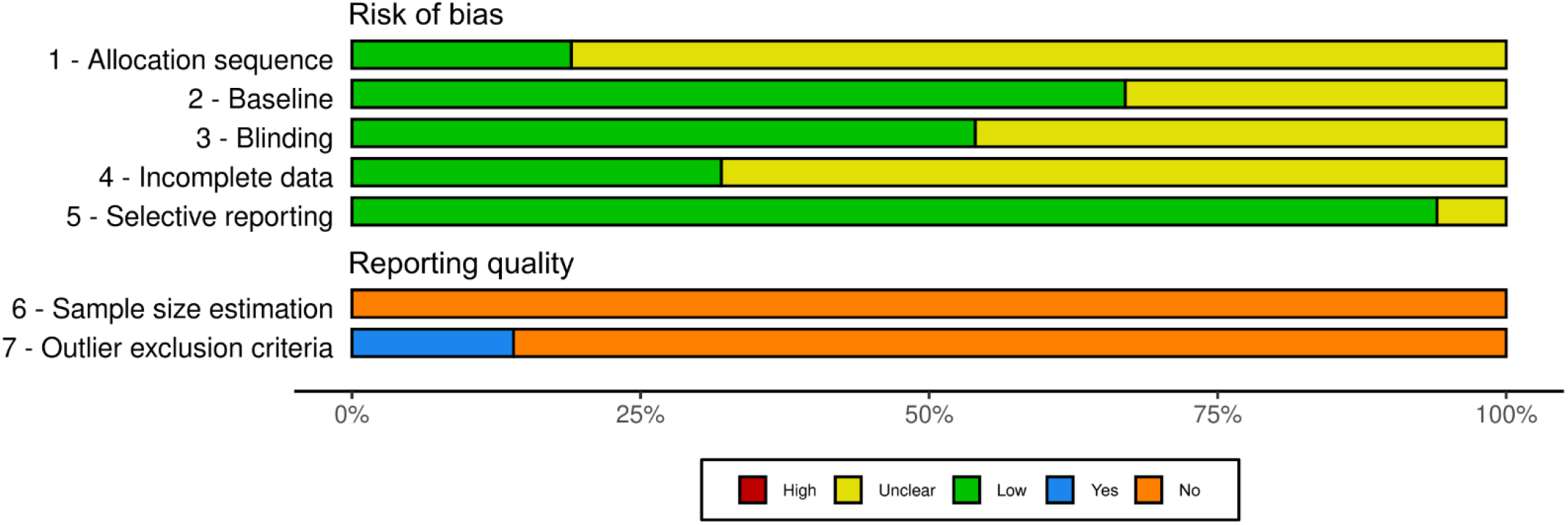
Risk of bias assessment of included studies. The risk of bias assessment was performed by two independent investigators based on the SYRCLE’s risk of bias assessment tool. Items 1 to 5 (Allocation sequence, Baseline, Blinding, Incomplete data and Selective reporting) account for methodological quality and were scored as presenting a high, unclear or low risk of bias. Sample size estimation and outlier exclusion criteria evaluate the reporting quality of the studies and were scored as “Yes” or “No”, meaning satisfactory or unsatisfactory reporting quality, respectively. Classification is given as the percentage of assessed studies (n *=* 100) presenting each score.

As for the reporting quality, more than 50% of the studies failed to report any information on the items assessed. Reporting quality was considered unsatisfactory when evaluating the report of inclusion/exclusion criteria, since there were no reports for this item in 86 (86%) of the studies. Sample size calculation was not reported in any of the studies. Individualized scores for each included study are available at https://osf.io/bgcdv.

## DISCUSSION

For the behavioral characterization of epileptic seizures and the discovery of new therapies to control epileptic seizures, different model organisms have been employed, including rodents, flies, worms, and fish [24]. Zebrafish has proved to be an efficient experimental model for chemically-induced epileptic seizures. The features of seizures are similar in humans: excessive neuronal discharges and progressive behavioral changes [7]. The response to pharmacological treatments is also evident in zebrafish. Larvae and adult animals that were pretreated with ASDs had seizures prevented when exposed to seizure-inducers [7,8,71,207]. Berghmans et al. (2007) [8], extended the studies by Baraban et al. (2005) [7] by testing 13 known ASDs against exposure to pentylenetetrazole (PTZ), validating the use of zebrafish in the search for new antiepileptic drugs and serving as a primary study, before being tested in other animal models, such as mammals [8]. The use of zebrafish in epilepsy research has grown exponentially in recent years, as this organism model offers many advantages: zebrafish have known homologs for 85% of the recognized epilepsy genes found in humans, has easy genetic manipulation, and can be maintained in large quantities, in addition to being used at different stages of development [32].

Here, we conducted a systematic review of studies that used chemical inducers to provoke seizures in zebrafish. For instance, most of the studies included in this review determined the behavioral responses and expression of genes involved in epileptogenesis to assess the mechanisms involved in epilepsy and discover new antiepileptic drugs, in addition to biomarkers of oxidative stress, neural activity and inflammation markers. The following subsections describe the effects of different chemical inducers of seizure and their capacity to reproduce behavioral phenotypes in zebrafish at different developmental stages.

### Pentylenetetrazole (PTZ)

In 180 studies that are included in this review, the seizure agent used is pentylenetetrazole (PTZ), a GABA_A_ receptor antagonist, that demonstrates characteristic and well-defined seizure behavior in zebrafish, both in larvae and in adults [7,9].

Baraban et al. 2005 characterized the behavior of epileptic seizures of larvae with 7 dpf using PTZ as a convulsant agent for 10 minutes and divided it into 3 stages: stage I — dramatically increased swimming activity, stage II — whirlpool swimming behavior, and stage III — clonus-like seizures followed by loss of posture, when the animal falls to one side and remains immobile for 1–3 s. Also, in this study the latency for stage I, II, or III seizure onset depended on PTZ concentration: at higher PTZ concentrations, the latency to a given seizure stage was shorter than that measured at lower concentrations. Another finding was that lower concentrations of PTZ (2.5 and 5.0 mM) evoked only Stages I and II of seizure behavior. Still, nearly 75% of all fish exhibited at least one clonus-like convulsion (Stage III) in the presence of 15 mM PTZ. In addition, the distance traveled also increased at the concentrations tested. This study serves as the basis for the following studies using behavioral characterization to study new drugs with antiepileptic activity [25,128,33].

In a study conducted by Afrikanova et al. (2013) [50], an alternative protocol was employed to investigate the effects of acute exposure in zebrafish larvae. Larvae of 6 dpf were placed in 96-well plates with ASD or vehicle for 18 hours of pre-treatment. After pre-treatment, 20 mM PTZ were added. Then animals were habituated for 5 min and locomotor activity was assessed for 30 min. The epileptic seizure stages observed were the same as in the study by Baraban et al. (2005) [7], but the difference between the two studies is the concentration of PTZ (15 mM PTZ, Baraban; 20 mM PTZ, Afrikanova) and the sequence of application of the drugs used (PTZ added before ASD, Baraban; ASD added before PTZ, Afrikanova). Other researchers began to use the protocol by Afrikanova et al, 2013 in their epilepsy experiments using zebrafish larvae (Barbalho, Lopes-Cendes, and Maurer-Morelli 2016; Brillatz, et al. 2020).

In adult zebrafish, the protocols followed the 3 stages characterized by Baraban et al. 2005 until the study by Mussulini et al. 2013 [9], which characterized 6 stages of epileptic seizures induced by PTZ where different concentrations of the inducer were evaluated. The stages were defined as: (0) short swim, (1) increased swimming activity and high frequency of opercular movement, (2) erratic movements, (3) circular movements, (4) clonic seizure-like behavior, (5) fall to the bottom of the tank and tonic seizure-like behavior, (6) death. The seizure stages were evaluated for 20 min, and they found that 10 mM was the ideal concentration to induce seizures, with animals reaching stage 5 in 240 s, approximately.

As we can see in Table 1, PTZ was used in 125 studies with zebrafish at larval stages. In larvae, the most common concentration was 20 mM for 30 min, where an increase in epileptic seizure stages and distance traveled was observed. At this concentration, PTZ increases gene expression of *c-fos* and *bdnf* [42,52,158]. The same was seen in other widely used concentrations, such as 10 mM and 15 mM [107,114,159]. The study by Lopes et al. (2016)[77] showed that approximately 24 hours after an acute exposure to 15 mM of PTZ for 60 min, *c-fos* gene expression was not statistically different when compared to the control group, indicating that this transcript factor, although altered soon after an acute induction, returns to its baseline levels after 1 day. In addition, changes in oxidative stress were observed, such as increased production of reactive oxygen species (ROS) and decreased activity and gene expression of antioxidant enzymes such as SOD, CAT and GPx, which may lead to neuronal death, at a concentration of 15 mM for 20 min in the study by Jin, et al. 2018. The expression of genes involved in neuroinflammation, such as il-1B and il-6, appear already increased at 5 mM PTZ exposure for 3 and 30 min [143].

In adults, the most common PTZ exposures were at concentrations of 7.5 and 10 mM for 10 to 20 minutes, where it was found that zebrafish showed increased distance traveled, speed, as well as increased gene expression of *c-fos* also seen in the larval stages [34,70]. Few studies evaluated *bdnf* expression, that is involved in neuroplasticity, in adults, and the majority used 170 mg/kg ip. Injection to induce seizures and some did not find any alteration (Choo et al., 2018, 2019; Jaiswal et al., 2020), while some observed a decrease in this gene expression [88,133]. Little is known about oxidative stress in adult zebrafish exposed to PTZ. Fontana et al. (2019) [129] found that 10 mM PTZ for 20 min causes an increase in thiobarbituric acid reactive species (TBARS) and carbonyl proteins, indicating lipid peroxidation, in addition to a decrease in non-protein thiol levels (NPSH).

PTZ was recently used chronically in an attempt to validate a kindling model in zebrafish [132,169], but in existing studies, seizures are evaluated shortly after induction with PTZ, with no spontaneous seizures in the animal. This is a gap that still needs to be investigated in zebrafish. Rodent *kindling* is a well-established chronic model that involves repeated electrical stimuli of sub effective intensity or repeated low-dose administration of a seizure-inducer until the induction of complete tonic-clonic seizures [220–222]. Kundap, Paudel et al. (2019) [132] tested PTZ 170 mg/kg i.p. for 10 days, while Kumari et al. (2020)[169] used 1.5 mM PTZ by immersion for 22 days and both studies found an increase in seizure scores over time, but without spontaneous crises. More studies are necessary to better develop a useful protocol to induce *kindling* in zebrafish to understand the epileptogenesis process and discover new treatments for epilepsy.

Most electrophysiological studies in zebrafish exposed to PTZ to record local field potential were performed in the larval phase, showing interictal and ictal-like epileptiform discharges with different concentrations of PTZ [116,122,128,136]. These results corroborate other studies carried out in mammals, where behavioral changes, alterations in the gene expression of *c-fos, bdnf*, and changes in oxidative stress are also observed [223–225]. Through this, PTZ in different concentrations seems to be a adequate chemical inducer of epileptic seizures in zebrafish used in most studies in this review. However, there are discrepancies in some results, making necessary to conduct more studies on these parameters and the dose/concentration should be decided according to the objectives of the study.

### Kainic acid (KA)

The second major chemical inducer of seizures found in this systematic review is KA, a potent excitatory neurotransmitter that is commonly used to induce seizures in animal studies [226,227]. The mechanism of action of KA in inducing seizures is through activation of the ionotropic glutamate receptor, specifically the kainate receptor subtype. Activation of these receptors leads to the influx of sodium ions into the neurons, which triggers a cascade of events leading to neuronal hyperexcitability and ultimately seizure activity [228]. The first study in this review that used kainic acid to induce seizures in zebrafish was conducted by Kim et al. 2010 [33], to determine whether PTZ- or KA-induced seizures influence cell proliferation in zebrafish larvae using bromodeoxyuridine (BrdU) to label dividing cells. They found that exposure to 200 µM KA for 10 min decreased the number of BrdU labeled cells in 5 dpf larvae in different brain areas, and the same was seen after exposure to 10 mM PTZ for 10 min, which indicated that seizures result in a massive reduction in cell proliferation in wide-ranging areas of the developing brain. In electrophysiology, KA at 50 μM increased epileptiform discharges characterized by short duration interictal events (100−200 ms) and long-lasting bursting discharges (4−5 s) occurring 8 times per minute on average. Menezes et al. (2014) [62] have shown that exposure of larvae to 100–500 μM KA decreased locomotor activity at 7 dpf and increased locomotion at 15 dpf. In addition, pre-exposure to KA at 24 hpf reduces the susceptibility of juvenile fish to generate seizures when later exposed to KA. Still in larvae, Feas, et al. 2017 [86] tested concentrations between 1.25 to 10 mM of KA for 25 min and an increase in spontaneous movements in 5 dpf larvae was seen.

Alfaro et al. (2011) [37] conducted an experiment using kainic acid (KA) to assess the behavior of adult zebrafish following intraperitoneal injections of varying doses (1, 2, 4, 6, and 8 mg/kg) of KA. The seizure stages used to characterize the seizures were similar to those observed with PTZ: stage I involved immobility and hyperventilation of the animal, stage II displayed whirlpool-like swimming behavior, stage III showed rapid movements from right to left, stage IV exhibited abnormal and spasmodic muscular contractions, stage V displayed rapid whole-body clonus like convulsions, stage VI involved sinking to the bottom of the tank and spasms for several minutes, and stage VII resulted in death. They observed a dose-dependent increase in seizure scores. Moreover, higher doses (above 6 mg/kg) significantly reduced the latency to the first stage V seizure compared to lower doses. Animals administered with 8 mg/kg of KA exhibited *status epilepticus,* defined as one continuous unremitting seizure lasting longer than 30 min. Sierra, et al. (2012) [47] also used 6 mg/kg of KA to induce seizures in adult zebrafish by intraperitoneal injection and an increase in seizure scores was seen in 60 min, with some animals reaching *status epilepticus*.

The study by Mussulini et al. (2018) [113] aimed to investigate the effects of kainic acid-induced *status epilepticus* on glutamate uptake and behavioral parameters in adult zebrafish. The authors found that kainic acid induced a significant decrease in glutamate uptake in the forebrain of zebrafish, suggesting that glutamate neurotransmission is altered during *status epilepticus*. Additionally, the study showed that zebrafish exhibited several behavioral changes following kainic acid administration, including hyperactivity, freezing, and erratic swimming patterns. The authors suggest that alterations in glutamate uptake and release in the forebrain may underlie these behavioral changes. Kundap, et al. 2019 [132] compared a single dose of KA 3 mg/kg with 10 days of PTZ 80 mg/kg and the results showed that KA increased seizure score at day 1 but decreased with time, while PTZ increased the seizure scores during the 10 days.

More recently, the study by Heylen et al. (2021) [196] investigated the use of pericardial injection of kainic acid to induce a chronic epileptic state in larval zebrafish. The authors found that this method resulted in a sustained increase in seizure frequency and duration over a period of several days, indicating the development of a chronic epileptic state. The study also identified changes in gene expression in the brain of zebrafish following KA injection, including upregulation of genes involved in inflammation and neurodegeneration at different time points after the microinjection. These results are accompanied by an increase in epileptiform discharges starting 2 days post-injection. In oxidative stress, Farias et al. (2022) [215] showed that 5 mg/kg increased the levels of reactive oxygen species (ROS) and lipid peroxidation, but did not alter the activity of antioxidant enzymes like SOD, CAT, glutathione reductase (GR) and glutathione peroxidase (GPx), just an increase in the SOD/CAT activity ratio. Overall, the studies reviewed highlight the usefulness of KA as a reliable chemical inducer of seizures in zebrafish models and provide insights into the underlying mechanisms and behavioral outcomes of KA-induced seizures.

### Pilocarpine (PILO)

Pilocarpine, a muscarinic cholinergic receptor agonist, is another agent used to induce epileptic seizures that progress to status epilepticus in rodents, and behavioral and biochemical changes are also observed when administered to adult zebrafish (Lin et al. 2017; Paudel et al. 2020).

Ten studies used pilocarpine as a seizure inducer in zebrafish. Vermoesen, et al. (2011) [41] tested PTZ and pilocarpine in *Tg(fli1a:EGFP)y1* zebrafish larvae (7dpf) and found an increase of total movement when 20 mM pilocarpine was administered by immersion for 10 minutes. Antiseizure effects of three antidepressants (citalopram, reboxetine, bupropion) against pilocarpine were tested, but only citalopram minimized the effect of pilocarpine. Lopes et al. (2016) [77] found that 60 mM pilocarpine administered for 60 min to larvae resulted in increased seizure activity, as evidenced by the rapid, rhythmic, and repetitive movements. The knockdown of carboxypeptidase A6 (*Cpa6*) using morpholino antisense oligonucleotides reduced the response to seizure-inducing drugs and caused changes in the expression levels of mRNAs encoding signaling molecules. Specifically, the expression of genes associated with neurotransmitter release, ion channel activity, and G protein-coupled receptor signaling was altered. Winter, et al. 2017 [95] showed electrophysiological outcomes in *elavl3:GCaMP6s* zebrafish larvae (4 dpf) exposed to 1 mM of pilocarpine for 55 minutes. This resulted in an increase in event power of neuronal network events (mV^2^), but not in frequency (Hz). The electrophysiology and the imaging data for pilocarpine suggested relatively widespread reduction in activity, however this was predominantly observed in the rhombencephalon. This was the only study that performed electrophysiological recording to evaluate the effect of pilocarpine in zebrafish.

Recently, the study by Paudel et al. (2020) [175] investigated the behavioral and biochemical changes induced by pilocarpine in adult zebrafish in a chronic seizure-like condition. The researchers administered pilocarpine (400 mg/kg) to the zebrafish through intraperitoneal injection and observed the development of seizure-like behaviors over time through the scores: 0 - Normal swimming, 1 - Jittery movement at the top of the tank, 2 - Ataxia or hyperactivity, 3 - Circular movement, circling around small area, 4 - Erratic burst movement with loss of posture/corkscrew swimming. The results showed that pilocarpine induced seizure-like behaviors in adult zebrafish, characterized by hyperactivity, seizures, and loss of posture. The chronic administration of pilocarpine also led to biochemical alterations, as significant increases in mRNA expression of *hmgb1, tlr4, il-1*; and decreased expression of *nfκb*. In addition, the group exposed to pilocarpine decrease GABA levels. The seizure behavior was also observed by Budaszewski et al., (2021) [184], where adult zebrafish were injected with pilocarpine (350 mg/Kg, i.p.) and 13% of animals showed lower scores (I–III) and 87% reached scores IV-V. Only fish presenting high scores (IV-V) were used in the PILO group to evaluate behavior on the light/dark test, open tank, aggressiveness test and biochemical parameters at different times. The study identified an increase in aggressive behavior and cell death in the PILO group, with increased levels of cleaved proteins caspase 3 and *parp1* 24 hours after seizure-like behavior induction. In addition, there were decreased protein levels of *psd95* and *snap25* and increased BrdU positive cells 3 days after induction. Persistent aggressive and anxiolytic-like behaviors were still detected 30 days after the induction, although most synaptic and cell death marker levels seemed normal.

Most recent studies with pilocarpine in this review [176,205] showed that a second hit with 80 mg/kg PTZ after 10 days of 400 mg/kg pilocarpine exposure increased the time spent in the upper half of the observation tank, and changed the expression of different genes, like increase in *hmgb1*, *tlr4*, *NF-kB*, *tnf-α*, *bdnf*, *creb1* and a decrease in *npy* expression. Paudel et al. (2021) [205] also observed an increase in *c-fos* expression. The use of pilocarpine as an epileptic seizure inducer in zebrafish models has provided important insights into the underlying neurochemical changes that occur during epilepsy and has implications for the development of new antiepileptic treatments.

### Picrotoxin (PTX)

The mechanism of action of picrotoxin (PTX) is similar to that of PTZ. PTX is a non-competitive GABA-A receptor antagonist, known to induce tonic and/or clonic seizures in different species, including rats and mice [229–231].

The first study to use picrotoxin in zebrafish was by Wong, et al. (2010) [35], where they tested PTZ, caffeine and PTX on behavioral and cortisol levels. The results showed that adult zebrafish exposed to 100 mg/L PTX for 20 minutes had an increase in bursts of hyperactivity, spasms, corkscrew swimming, circular swimming and time spent in the upper half of the tank; cortisol levels were increased. Baxendale, et al. (2012) [42] found an increase in *fos* expression in zebrafish embryos exposed to 300 μM PTX and the same was seen with 20 mM of PTZ.

The studies using zebrafish with PTX as a seizure-inducer basically evaluated behavioral outcomes and found alterations in distance traveled, average of movements and speed [14,117,148,199]. There are discrepancies in locomotor parameters such as distance traveled and velocity between the studies, and it could be due to the use of different doses or concentrations tested and the time of exposure. In electrophysiology, just one study by Bandara et al. (2020) [148] showed an increase in epileptiform-like discharges and burst frequencies. In short, further studies are needed to elucidate and characterize epileptic seizures and molecular alterations caused by picrotoxin in zebrafish.

### Other chemical seizure-inducers

There are other drugs that are used to induce seizures in zebrafish, but they are not as usual, including caffeine, domoic acid, allylglycine, ethyl ketopentenoate, ginkgotoxin (GT), strychnine, 4-aminopyridine (4-AP), and 1,3,5-Trinitroperhydro-1,3,5-triazine (RDX), among others.

Domoic acid is a neurotoxin produced by certain species of algae that can cause seizures and other neurological symptoms in animals that consume contaminated shellfish [232]. Studies included in this review showed that domoic acid increases the susceptibility to seizures when administered in larvae with other drugs like PTZ and contaminants [26,30,31]. Lefebvre, et al. (2009) [28] showed that an intraperitoneal injection of 0.47 µg/g and 1.2 µg/g of domoic acid in adult zebrafish increased abnormal behavior and mortality.

Another drug used to study epileptic seizures in zebrafish is 4-amynopiridine (4-AP), a potent voltage-gated potassium channel blocker [233]. 4-AP seems to cause seizure behavior in zebrafish [27,83], increase epileptiform events (4-AP 4mM) [136], and hyperexcitability in the hindbrain or lateral optic tectum [168]. Also, Ellis et al., (2012) [43] showed an increase in c-fos expression in a concentration dependent manner at concentrations between 0.6 to 2.5 mM for ten minutes.

Caffeine is a central nervous system stimulant that has been shown to induce seizures in zebrafish at high concentrations [94]. Also, Wong et al. (2010) [35] notice that 250 mg/L caffeine for 20 minutes alters zebrafish locomotor behavior, increasing spasms and bursts of hyperactivity, but decreasing distance traveled. Another study by Wong, et al. (2010b) showed that 15 minutes of exposure to caffeine 100 mg/L for 2 weeks increased erratic movements in the animal.

Leclercq, et al., (2015) [13] tested concentrations of allylglycine (AG) (30 to 300mM) for 480 minutes in 7 dpf zebrafish larvae and found an increase in swimming activity and a whole body seizure. Also, 300 mM AG for 120 minutes presents epileptiform discharges in electrophysiological analyses and a decrease of GABA in brain homogenates. Ethyl ketopentanoate (EKP) is a derivative of AG and both are inhibitors of glutamate decarboxylase (GAD). Zhang, et al., (2017) [96] tested different concentrations of EKP (200-800 µM), and the results showed that EKP evoked robust convulsive locomotor activities, excessive epileptiform discharges and upregulated *c-fos* expression in zebrafish. Li, et al. (2020) [173] tested the concentration of 1 mM and the results also showed hyperlocomotion and epileptiform discharges. Alterations in locomotor behavior and epileptiform activity also were seen by Sourbron, et al. (2018) [144].

The other drugs that appear in Table 1 also showed capacity to alter the zebrafish behavior, like 0.2 to 1 mM of ginkgotoxin (GT) that increases seizure score and distance traveled [45], 1 mM RDX and NMDA (different concentrations), that increase average seizure score [48,174], and 4 µM tetramethylenedisulfotetramine (TETS) that increases distance traveled and causes seizure-like behavior [203]. Despite these results, further studies are needed to better characterize epileptic seizures in zebrafish and their molecular and electrophysiological changes.

### Risk of bias and reporting quality

To assess risk of bias and reporting quality we randomly chose a sample of 100 articles included in this review. The majority of studies were rated as having a low risk of bias for selective reporting, suggesting that most studies included in the review reported all mentioned outcomes. This, however, should be interpreted with caution as protocol preregistration is not standard in preclinical literature, and we thus cannot rule out that other outcomes were collected and not reported. In addition, the results indicate that more than half of the studies have a low risk of bias for baseline characteristics of animals and blinding, but these points still could be potential biases in some studies. The fact that randomization procedures and incomplete data were frequently rated as unclear is also a concern, as this suggests that some studies have not adequately reported on these important aspects of study design. The lack of information might compromise the credibility and reproducibility of data. This information is necessary to assess the methodological rigor of the studies and for other researchers to repeat experimental procedures [234,235].

As for reporting quality, the fact that more than 50% of studies failed to report information on the items is concerning. The lack of reporting on inclusion/exclusion criteria in the majority of studies is particularly concerning, as this makes it difficult to assess the validity of study results. Additionally, the fact that sample size calculation was not reported in any of the studies is another important limitation, as it makes it difficult to assess the statistical power of the studies.

## Conclusions

This systematic review carried out a detailed literature search, looking for studies on the effects of chemical seizure inducers in zebrafish. We summarized the information to provide a tool for the researcher to select the best protocol to chemically-induce seizures in zebrafish larvae and adults to study epileptic crisis or assess antiseizure effects of different compounds. Choosing the best model needs to consider specific objectives to address the research question and outcomes of interest. Although there are several available inducers for epileptic seizures, studies using PTZ are by far the most common and extensively characterized. However, it is crucial to consider the risk of bias and quality of reports observed in the studies to improve the protocol to be used.

It is clear that seizure models are continually being improved; however, better standardization and better models, such as kindling, are still needed to advance knowledge in the field of epilepsy.

## AUTHOR CONTRIBUTION

**Rafael Chitolina:** conceptualization, data curation, formal analysis, investigation, methodology, project administration, visualization and writing - original draft; **Matheus Gallas-Lopes**: conceptualization, data curation, investigation, methodology, visualization and writing - review & editing; **Carlos Guilherme Rosa Reis**: conceptualization, investigation, methodology, visualization and writing - review & editing; **Radharani Benvenutti**: conceptualization, investigation, methodology, visualization and writing – review & editing; **Thailana Stahlhofer-Buss**: investigation, visualization and writing – review & editing; **Maria Elisa Calcagnotto**: conceptualization, investigation, methodology, project administration, supervision, visualization and writing – review & editing; **Ana P. Herrmann**: conceptualization, investigation, methodology, project administration, supervision, visualization and writing – review & editing; **Angelo Piato:** conceptualization, investigation, methodology, project administration, supervision, visualization and writing – review & editing.

## CONFLICT OF INTEREST

The authors declare no conflict of interest.

## ACKNOWLEDGMENTS

We thank the Conselho Nacional de Desenvolvimento Científico e Tecnológico (CNPq, proc. 303343/2020-6), Coordenação de Aperfeiçoamento de Pessoal de Nível Superior - Brasil (CAPES), and Pró-Reitoria de Pesquisa (PROPESQ) at Universidade Federal do Rio Grande do Sul (UFRGS) for funding and support.

## FUNDING

RC is recipient of a fellowship from CAPES.

## Notes

### Competing Interest Statement

The authors have declared no competing interest.

